# Epigenetically constrained astrocyte states underlie prefrontal cortex vulnerability in Down syndrome–associated Alzheimer’s disease

**DOI:** 10.64898/2026.04.17.719050

**Authors:** Chuhanwen Sun, Raina Thomas, Cherie Stringer, Kyriakitsa Galani, Li-Lun Ho, Na Sun, Ashley Renfro, Sierra Wright, Rosalind Firenze, Li-Huei Tsai, Elizabeth Head, Manolis Kellis, Jiekun Yang

## Abstract

Down syndrome (DS), caused by trisomy 21, confers a near-universal risk for Alzheimer’s disease (AD), yet individuals exhibit marked variability in cognitive decline, suggesting the presence of cellular mechanisms that modulate vulnerability and resilience. However, these mechanisms remain poorly defined in the human brain. Here, we integrate matched single-nucleus RNA-seq and ATAC-seq profiles from the prefrontal cortex (PFC) and amygdala (AMY) of age-matched individuals with DS with and without AD (DSAD), enabling direct comparison within a shared genetic background.

We identify basal astrocytes in the PFC as a selectively vulnerable cell state in DSAD, characterized by both reduced abundance and coordinated transcriptional and regulatory reprogramming. This state exhibits a shift away from homeostatic support functions, with decreased cytokine signaling and lipid-handling programs, alongside increased steroid- and nuclear receptor–associated activity. Concomitantly, chromatin accessibility profiling reveals reduced engagement of immune- and stress-responsive transcription factor programs, including AP-1, STAT, and BACH families, with linked regulatory perturbations at loci such as ABCA1, DAB2IP, and IL1RAP.

Together, these findings define a previously unrecognized astrocyte state marked by epigenetic constraint and diminished responsiveness to stress and inflammatory signals, distinguishing it from classical reactive astrocyte phenotypes. Our results nominate PFC basal astrocytes as a key locus of vulnerability in DSAD and suggest that failure to mount appropriate astrocyte responses, rather than overt activation alone, may contribute to neurodegenerative progression.

## Introduction

Down syndrome (DS), a genetic condition that is most often linked to trisomy 21, is characterized by lifelong cognitive impairments and the gradual accumulation of neuropathological features of Alzheimer’s disease (AD)^1,2^. By the age of 40 years, nearly all individuals with DS present with amyloid plaques and tau neurofibrillary tangles. With aging, they develop an aggressive, genetic form of AD that results in cognitive decline and carries a lifetime risk of over 90%. Although the burden of AD is high for people with DS, there are limited clinical trials and no approved treatments for this population^1^. A deeper mechanistic understanding of the unique etiology of the AD that develops in Down syndrome, or ‘DSAD’, is needed for the development of targeted and safe therapies. Single-nuclei RNA- and ATAC-sequencing have been powerful tools for generating a cell-specific understanding of the transcriptomic and epigenetic states in the human brain that underlie AD^3–8^. Past studies have utilized these tools to identify cell-specific changes that may contribute to cognitive impairments in mouse models of DS, including the enrichment of a senescence-associated transcriptional signature in oligodendrocyte precursor cells (OPCs) and a greater proportion of microglia in an intermediate state of reactivity in the Ts65Dn model of DS^9–11^. However, there is limited information at the single-cell level for changes in humans with DS or DSAD. In one snRNA-seq study of 9 DS brains versus 20 non-DS controls, Palmer et al. reported that the greatest gene expression changes were found in microglia^12^. Their analysis indicated an aged microglial state with transcriptomic hallmarks of activation and a loss of homeostatic gene expression in microglia in all DS samples, regardless of age, producing an aged microglial state in young (≤36 years) DS brains. More recently, Miyoshi et al. generated single-nucleus transcriptomes from postmortem human brain samples from donors with DSAD and compared these to samples from patients with sporadic AD (sAD) and cognitively healthy controls^13^. Their work for the first time revealed regional and cell-type specific gene changes in DSAD that were shared with but also distinct from sAD. Due to the rarity of samples, Miyoshi et al. were not able to compare DSAD samples to those from persons with DS but without AD. This comparison could provide clues towards understanding resilience to AD in individuals with DS who remain cognitively stable despite neuropathology. Here, we utilize both snRNA-seq and snATAC-seq to study and compare the cellular diversity in the prefrontal cortex (PFC) and amygdala (AMY) from three groups: patients with DSAD, persons with DS but without AD and healthy, non-DS controls. Altogether, our results uncover brain region-specific and cell subtype-specific differences in DSAD, including transcriptional and epigenetic dysregulation in basal astrocytes of the PFC.

## Results

### Unveiling Cellular Diversity: Subtype-Specific Insights Across Brain Regions in Down Syndrome

We obtained frozen brain samples from the University of California Irvine, focusing on two regions: the prefrontal cortex (PFC) and amygdala (AMY). The study compared Down syndrome cases with (DSAD) and without (DS) Alzheimer’s disease. We profiled 24 samples using snRNA-seq and snATAC-seq, including 9 DSAD and 3 DS cases (**Fig. 1a**). Both groups had a 2:1 female-to-male ratio, with no significant differences in age or post-mortem interval (PMI; **Table 1, Supp. Fig. 1a**). Notably, the DS group had 2 APOE ε2 carriers and no APOE ε4 carriers, while the DSAD group had 4 APOE ε4 carriers and 1 APOE ε2 carrier, consistent with known risk profiles (**Supp. Fig. 1b**). All the DS individuals had a tangle stage of 3 compared to 6 for all the DSAD individuals.

**Figure 1.**
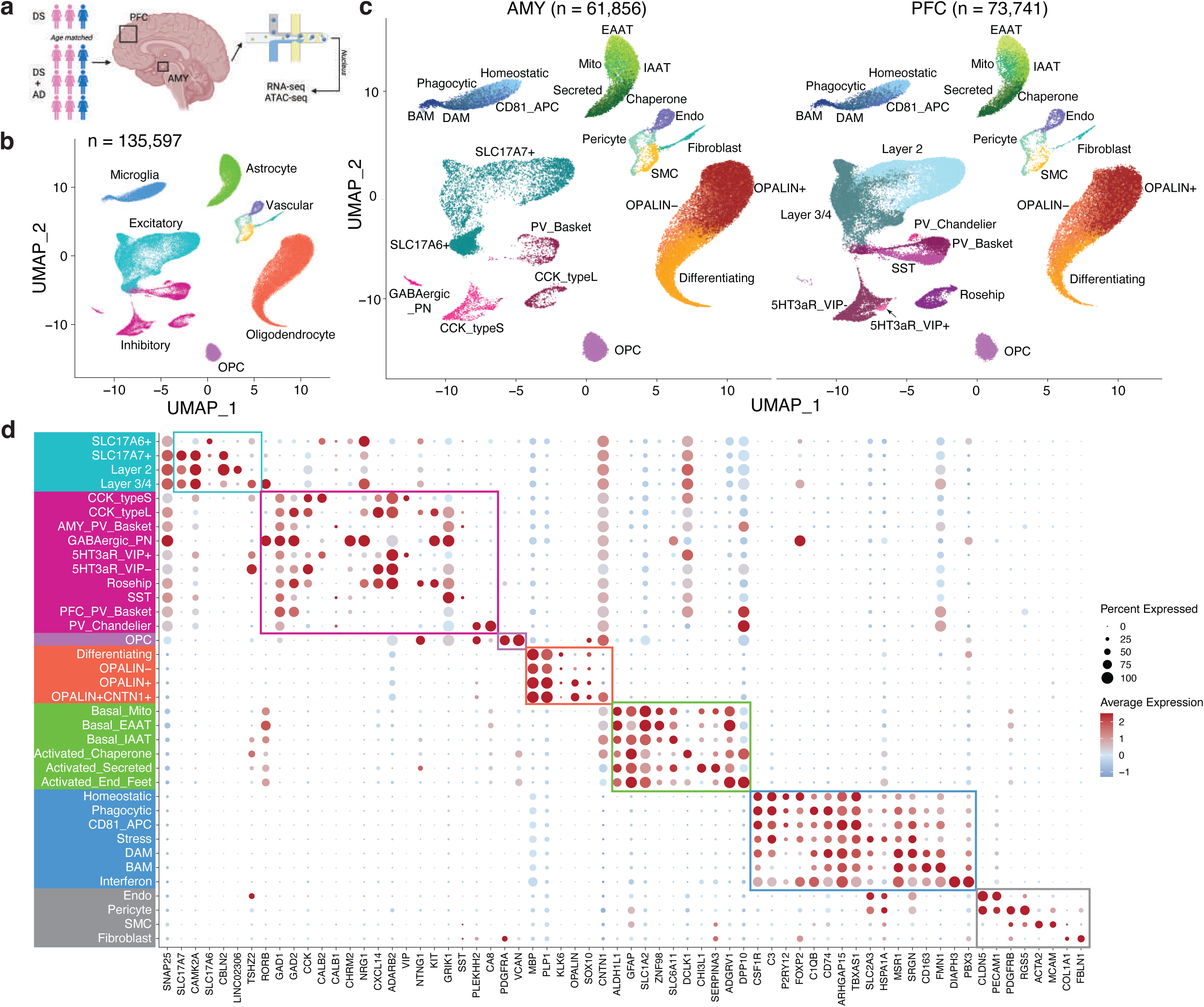
Single-cell profiling of the Down syndrome prefrontal cortex and amygdala with or without Alzheimer’s disease. a, Study design and cohort overview for matched single-nucleus RNA-seq and ATAC-seq profiling of the prefrontal cortex (PFC) and amygdala (AMY) from 12 individuals with Down syndrome, including 3 individuals with Down syndrome without Alzheimer’s disease (DS) and 9 individuals with Down syndrome-associated Alzheimer’s disease (DSAD), yielding 24 region-matched samples in total. b, Uniform manifold approximation and projection (UMAP) embedding of the integrated snRNA-seq atlas showing the major cell classes across all nuclei (n = 135,597). c, UMAP embeddings split by brain region showing annotated cell subtypes and states in AMY and PFC. d, Dot plot of selected marker genes across annotated subtypes and states. Dot size indicates the fraction of cells expressing each gene, and color indicates scaled average expression.

**Table 1.** Clinical information of patients. PMI, post-mortem interval; DS, Down Syndrome; AD, Alzheimer’s Disease.

| Patient ID | Age | Sex | PMI | APOE | Diagnosis | Tangle Stage | Plaque Stage |
| --- | --- | --- | --- | --- | --- | --- | --- |
| 1302 | 46 | M | 6.37 | 2/3 | DSAD | 6 | Stage C |
| 3108 | 49 | M | 2.2 | 3/3 | DSAD | 6 | Stage C |
| 3304 | 50 | F | 5 | 3/4 | DSAD | 6 | Stage C |
| 4815 | 55 | F | 4.58 | 3/4 | DSAD | 6 | Stage C |
| 3208 | 42 | F | 5 | 3/4 | DSAD | 5 | Stage C |
| 2906 | 45 | F | 2.75 | 3/3 | DSAD | 6 | Stage C |
| 515 | 51 | F | 2.72 | 2/3 | DS | 3 | Stage C |
| 3107 | 52 | F | 4.37 | 2/4 | DSAD | 6 | Stage C |
| 3317 | 62 | F | 7.03 | 3/3 | DS | 3 | Stage B |
| 3110 | 62 | F | 2.42 | 3/3 | DSAD | 6 | Stage C |
| 416 | 70 | M | 3.92 | 3/3 | DSAD | 6 | Stage B |
| 2715 | 72 | M | 4.87 | 2/3 | DS | 3 | None |

We confirmed chromosome 21 trisomy in all samples by inferring copy number variation (CNV) from snRNA-seq data (**Supp. Fig. 1c**)^14^. AMY samples generally showed more CNVs than PFC samples, potentially due to increased pathology in the AMY, leading to reduced sample quality or large-scale gene expression changes rather than true DNA alterations. We also detected partial trisomy in the PFC sample from individual 2715, consistent with previous findings^15^. Notably, the AMY sample from the same individual exhibited full chromosome 21 trisomy, suggesting a limitation of this approach, as this case is a confirmed partial trisomy without mosaicism (**Supp. Fig. 1c**)^15^.

We integrated data from the 24 snRNA-seq samples, applied dimensionality reduction, and visualized the 2D cell embeddings. Across 135,597 cells, we identified 10 cell types: excitatory and inhibitory neurons, oligodendrocyte progenitor cells (OPCs), oligodendrocytes, astrocytes, microglia, and vascular-related cells, including endothelial cells, pericytes, smooth muscle cells, and fibroblasts (**Fig. 1b,c**). The integration was consistent across sex, diagnosis, APOE status, brain region, age, PMI, and sample (**Extended Data Fig. 1a,b**). Mitochondrial read percentages were low due to nuclei isolation. As expected, neurons had more detected molecules and genes compared to other cell types (**Extended Data Fig. 1c**).

We sub-clustered each cell type and identified subtypes/states for 5 of the 10 cell types. Neuron subtypes were annotated according to standard naming conventions for each brain region. Excitatory neurons included layer 2 and layer 3/4 neurons in the PFC, and SLC17A6+ and SLC17A7+ neurons in the AMY (**Extended Data Fig. 1d**). Inhibitory neurons comprised several subtypes: 5HT3aR+ VIP+, 5HT3aR+ VIP-, Rosehip, somatostatin (SST), parvalbumin (PV) basket, and PV Chandelier in the PFC, along with CCK type S, CCK type L, PV basket, and GABAergic projection neurons in the AMY. Notably, GABAergic projection neurons were only observed in the AMY (**Fig. 1c**). AMY CCK type L neurons overlapped with PFC Rosehip neurons, and AMY CCK type S neurons overlapped with PFC 5HT3aR+ neurons, consistent with previous findings on euploid samples^16^.

We identified 4 oligodendrocyte subtypes: differentiating, OPALIN-, OPALIN+, and OPALIN+ CNTN1+ oligodendrocytes (**Extended Data Fig. 1e**). Astrocytes were classified into 6 subtypes—3 basal (mitochondria high, EAAT [excitatory amino acid transporter] high, IAAT [inhibitory amino acid transporter] high) and 3 activated (chaperone protein high, secreted protein high, and end feet; **Extended Data Fig. 1f**). Microglia were categorized into 7 subtypes: homeostatic, phagocytic, CD81+ antigen-presenting, stressed, disease-associated microglia (DAM), and brain-associated macrophages (BAM, **Extended Data Fig. 1g**). Canonical markers showed distinct expression patterns for the 30 identified subtypes/states (**Fig. 1d**).

### Validating Novel Astrocyte States Across Down Syndrome and Alzheimer’s Disease

To verify the consistency of cell types identified in DS and matched brain regions from euploid individuals, we integrated our snRNA-seq data with a publicly available dataset (**Extended Data Fig. 2a**)^16^. We observed strong consistency across variables such as sex, brain region, diagnosis, sample, and cell type. Our primary focus was the 6 *de novo* astrocyte states, which we successfully validated in the euploid samples (**Fig. 2a**). CytoTRACE analysis^17^ revealed a continuous progression from the three basal astrocyte states to three transcriptionally distinct non-basal states, here referred to as activated states, indicating a continuum of astrocyte functional programs rather than discrete, disease-specific activation (**Fig. 2b**).

**Figure 2.**
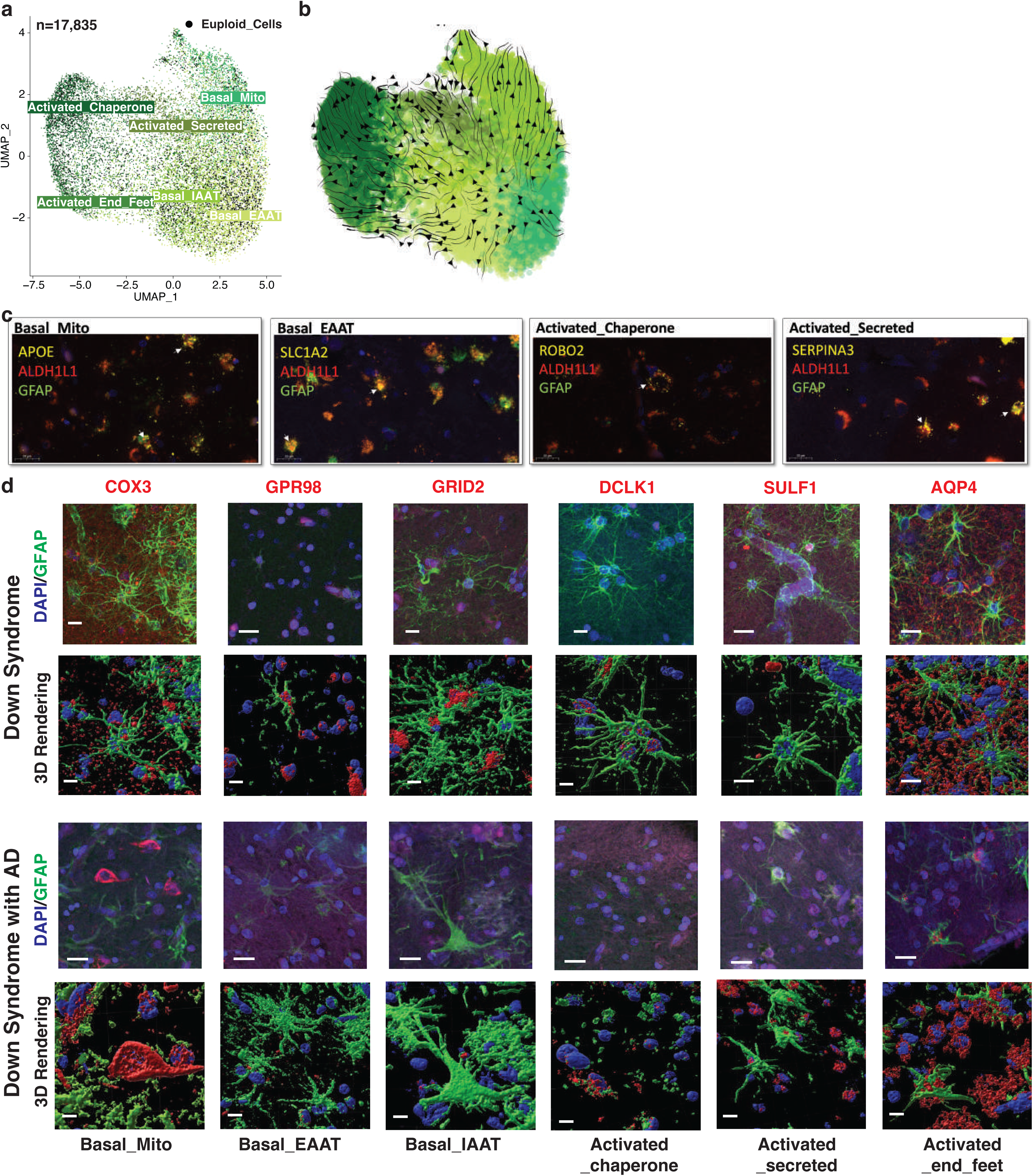
Six astrocyte states and their orthogonal validation in DS and DSAD. a, UMAP embedding of astrocytes (n = 17,838) showing six astrocyte states, including three basal states (basal mitochondria-high, basal EAAT-high, and basal IAAT-high) and three activated states (activated chaperone-high, activated secreted protein-high, and activated end-feet). b, Trajectory analysis of astrocytes arranged along pseudotime across basal and activated states. c, Representative RNAscope images showing co-detection of ALDH1L1, GFAP, and selected state-associated transcripts, including APOE for basal mitochondria-high astrocytes, SLC1A2 for basal EAAT-high astrocytes, ROBO2 for activated chaperone-high astrocytes, and SERPINA3 for activated secreted protein-high astrocytes. d, Representative immunofluorescence images and corresponding 3D renderings showing protein markers associated with the six astrocyte states in DS and DSAD tissue, including COX3, GPR98, GRID2, DCLK1, SULF1, and AQP4. Scale bars are indicated in the figure.

We further validated the six astrocyte states at both RNA and protein levels. Using RNAscope on selected samples, we observed co-localization of astrocyte markers (ALDH1L1 and GFAP) with specific markers for each state: APOE for the basal mitochondria-high state, SLC1A2 for the basal EAAT-high state, KCND2 for the basal IAAT-high state, ROBO2 for the activated chaperone-high state, SERPINA3 for the activated secreted protein-high state, and DPP10 for the activated end-feet state (**Fig. 2c, Extended Data Fig. 2b**).

Protein staining, particularly with 3D rendering, confirmed the presence of these six astrocyte states in both DS and DSAD individuals through co-staining of GFAP and state-specific markers (COX3, GPR98, GRID2, DCLK1, SULF1, and AQP4; **Fig. 2d**). Notably, AQP4 enrichment in this state is consistent with perivascular endfoot localization, suggesting that this population reflects astrocytes with prominent endfoot-associated features that may overlap with transcriptional programs labeled here as “activated,” rather than representing a purely activation-defined state^18^.

### Region-Specific Vulnerability of Astrocyte and Endothelial Cell in Alzheimer’s Within Down Syndrome

We analyzed cell type proportions across individuals and found strong correlations among astrocytes, microglia, and vascular-related cells in both the AMY and PFC (**Fig. 3a,b**). At the subtype level, these correlations remained strong, with some inhibitory neuron subtypes also correlating with them in the AMY, while OPCs and OPALIN+CNTN1+ oligodendrocytes correlated with them in the PFC (**Extended Data Fig. 3a,b**). Excitatory and inhibitory neurons also showed a strong correlation in terms of abundance in the PFC (**Fig. 3b, Extended Data Fig. 3b**).

**Figure 3.**
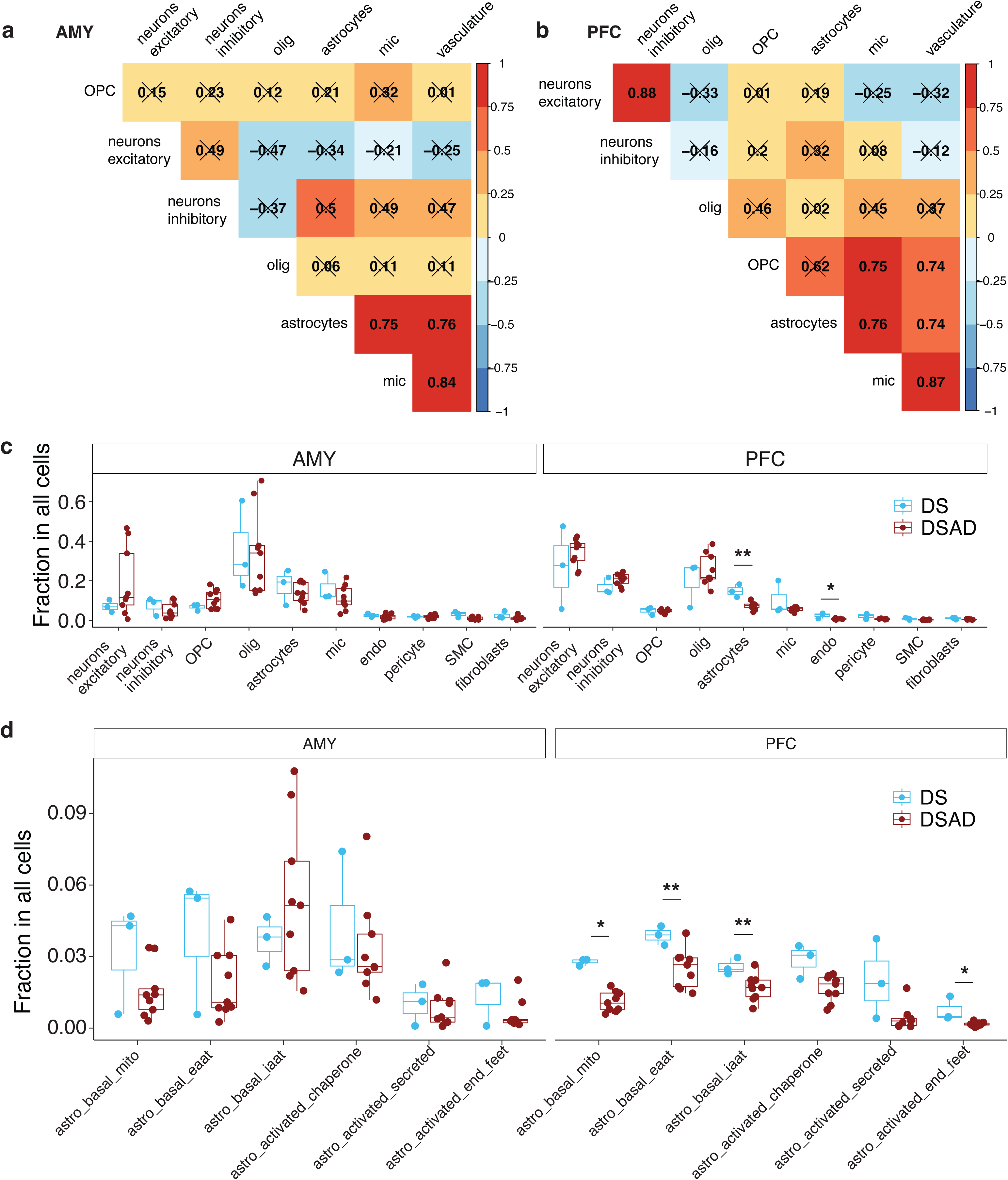
Region-specific cell-type and astrocyte-state proportion differences between DS and DSAD. a,b, Sample-level correlation heatmaps of cell-type abundances across donors in AMY (a) and PFC (b). Pearson correlation coefficient is indicated in each tile. c, Donor-level cell-type proportions in AMY and PFC comparing DS and DSAD. d, Donor-level astrocyte-state proportions in AMY and PFC comparing DS and DSAD. Boxplots summarize donor-level distributions and points denote individual donors. The lower and upper hinges correspond to the first and third quartiles. The upper- and lowerwhisker extends from the hinge to the largest and lowest value no further than 1.5 * IQR from the hinge. Wilcoxon Rank Sum tests were performed between DS and DSAD within each brain region and statistical significance are indicated in the figure. *P_≤_0.05; **P_≤_0.01.

To identify cell types most affected in DS and DSAD, we compared the proportions of cell types and subtypes across euploid, DS, and DSAD samples. The euploid samples, aged 40 to 69, had a similar age range to the DS and DSAD groups. Vascular-related cells were the only cell type with significant proportion differences between DS and euploid individuals, with a consistent increase in DS across both brain regions (**Extended Data Fig. 3c**). This differs from earlier reports of reduced endothelial progenitor cells in younger DS cohorts^19^, although differences in cohort composition and sample size may contribute to this discrepancy. In addition, DS samples exhibited a lower inhibitory-to-excitatory neuron ratio in both regions compared to euploid samples (**Extended Data Fig. 3d**), in contrast to previous findings in an independent DS cohort spanning a broader age range^12^. Given the limited sample size and differences in cohort structure, these observations should be interpreted with caution, and the sources of these discrepancies will require further investigation in larger and more systematically matched studies.

Comparing DS and DSAD samples, we observed a significant reduction in astrocytes and endothelial cells in the PFC, but not in the AMY, indicating a region-specific vulnerability associated with disease progression (**Fig. 3c**). No significant differences were detected in excitatory-to-inhibitory neuron ratios or OPC-to-oligodendrocyte ratios (**Extended Data Fig. 3e,f**). Within astrocytes, this decrease was primarily driven by loss of the three basal subtypes in the PFC, with the endfoot-associated subtype also reduced and the remaining activated subtypes showing a consistent downward trend (**Fig. 3d, Extended Data Fig. 3g**).

### Transcriptional Dysregulation in Basal Astrocytes in Down Syndrome and Alzheimer’s Disease

To identify cell-type-specific differentially expressed genes (DEGs) between DS and DSAD, we applied five statistical models to account for the limited sample size in the DS group. Covariates included in at least one model were age, sex, PMI, and APOE status. We used two algorithms: NEBULA, a negative binomial mixture model developed in-house^20^, and MAST, a cell-based model^21^. NEBULA, with various covariates, identified more DEGs in neurons, while MAST, without covariates, found more DEGs in astrocytes, microglia, and vascular cells (**Fig. 4a**). This suggests that transcriptional changes in neurons, particularly excitatory and inhibitory neurons in the PFC, are largely covariate-independent, whereas glial cells, but not basal astrocytes in the PFC, show covariate-dependent effects. OPCs and oligodendrocytes had a comparable number of DEGs across mixture and cell-based models.

**Figure 4.**
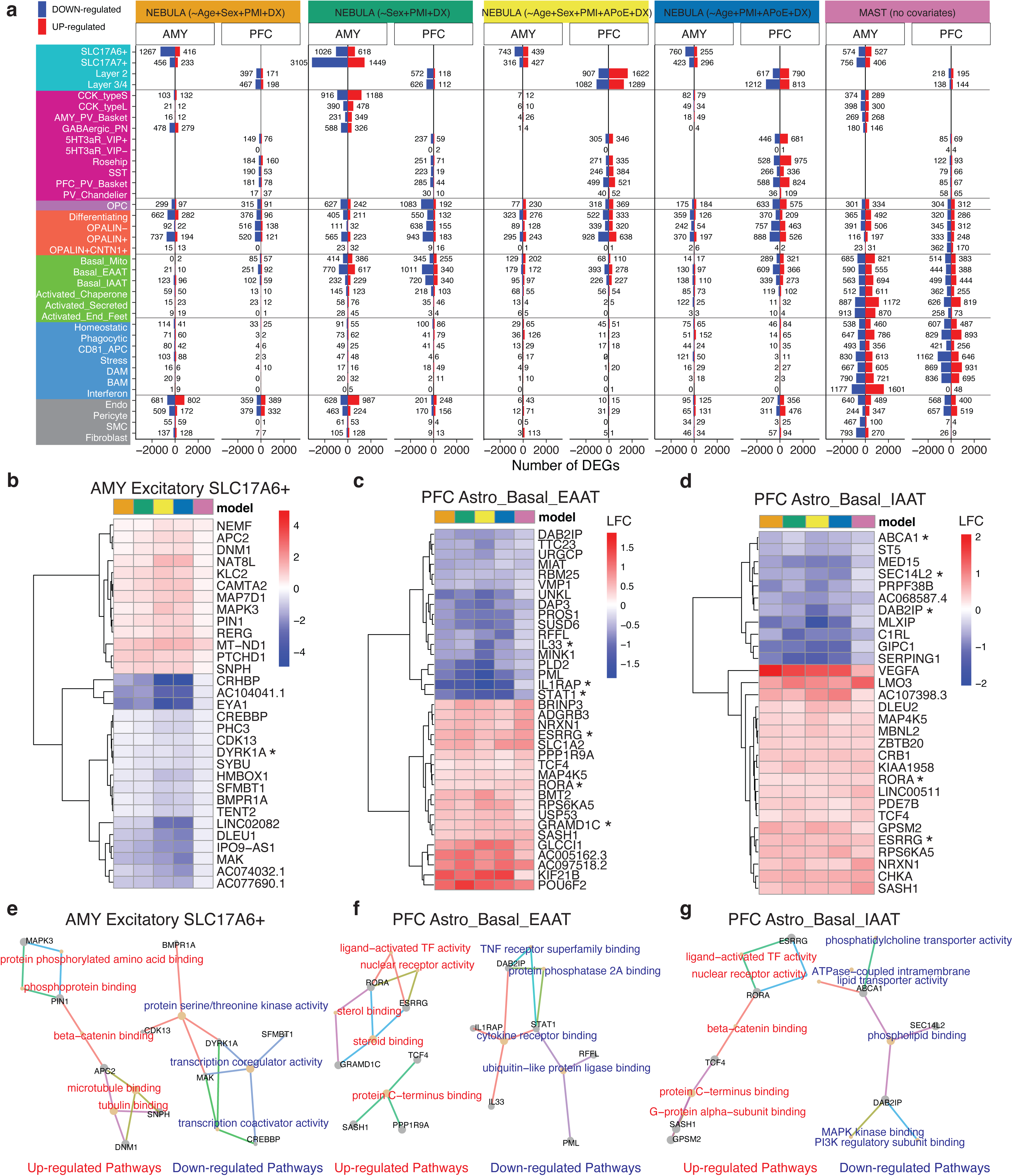
Consensus RNA dysregulation across statistical models highlights vulnerable PFC basal astrocytes. a, Summary of differentially expressed genes (DEGs) identified between DS and DSAD across five models, including four NEBULA models with different covariate combinations and one MAST model without covariates, stratified by brain region and cell subtype/state. Bars to the left and right of zero indicate genes decreased and increased in DSAD, respectively. b–d, Heatmaps of consensus DEGs in AMY SLC17A6+ excitatory neurons (b), PFC basal EAAT astrocytes (c), and PFC basal IAAT astrocytes (d). e-g, Network summaries of enriched gene-set terms for AMY SLC17A6+ excitatory neurons (e), PFC basal EAAT astrocytes (f), and PFC basal IAAT astrocytes (g). Each node represents a pathway or a gene. Up-regulated pathways are colored in red, down-regulated are colored in blue. Genes are linked to the pathways they belong to via edges, which are colored by pathways.

To prioritize robust signals, we focused on differentially expressed genes (DEGs) that were consistently detected across five models and exhibited concordant effect directions. This approach identified 365 consensus DEGs across specific cell subtypes and brain regions. For example, PICALM was downregulated in CD81D antigen-presenting microglia in the AMY in DSAD, consistent with its established role in AD^22^, and DYRK1A was reduced in SLC17A6+ excitatory neurons in AMY (**Fig. 4b**). Cell types exhibiting the largest number of consensus DEGs included SLC17A6D excitatory neurons in the AMY, basal EAAT and IAAT astrocytes in the PFC, OPCs in the PFC, and multiple oligodendrocyte subtypes (differentiating, OPALIND, and OPALIND) in the PFC (**Fig. 4b-d, Extended Data Fig. 4a-d**).

Although the number of consensus DEGs was modest, pathway-level analysis revealed coherent biological programs distinguishing DSAD from DS (**Fig. 4e-g, Extended Data Fig. 4e-g**). In basal EAAT astrocytes in the PFC, genes associated with steroid and nuclear receptor signaling, including ESRRG, RORA, and GRAMD1C, were upregulated in DSAD, whereas cytokine receptor–associated genes such as IL33, IL1RAP, and STAT1 were downregulated (**Fig. 4f**). A similar pattern was observed in basal IAAT astrocytes, with increased expression of ESRRG and RORA and concurrent downregulation of phospholipid-handling genes, including ABCA1, SEC14L2, and DAB2IP (**Fig. 4g**).

Ligand–receptor analysis revealed reduced JAG1–NOTCH2 signaling between astrocytes and pericytes/smooth muscle cells in both the AMY and PFC (**Extended Data Fig. 4h,i**). Given the role of NOTCH signaling in glial–vascular communication^23^, this reduction is consistent with the coordinated depletion of astrocyte and vascular populations and suggests disruption of astrocyte–vascular interactions in DSAD.

### snATAC-seq Recapitulates Astrocyte Heterogeneity and Confirms Their Vulnerability in DSAD

We generated snATAC-seq profiles from the same 24 samples and applied standard quality control based on transcription start site (TSS) enrichment and unique fragment counts (**Extended Data Fig. 5a-d**). After filtering, quality metrics were comparable across samples, and cells integrated consistently across diagnosis, brain region, and individual samples (**Extended Data Fig. 5e**). Clusters were annotated using gene activity scores for canonical cell type markers (**Fig. 5a,b**). Among 144,797 high-quality cells, we identified the same major cell types as in the snRNA-seq data, including excitatory and inhibitory neurons, OPCs, oligodendrocytes, astrocytes, microglia, and vascular-related cells. Consistent with the transcriptomic analysis, astrocytes showed a significant reduction in the PFC in DSAD relative to DS (**Extended Data Fig. 5f**).

**Figure 5.**
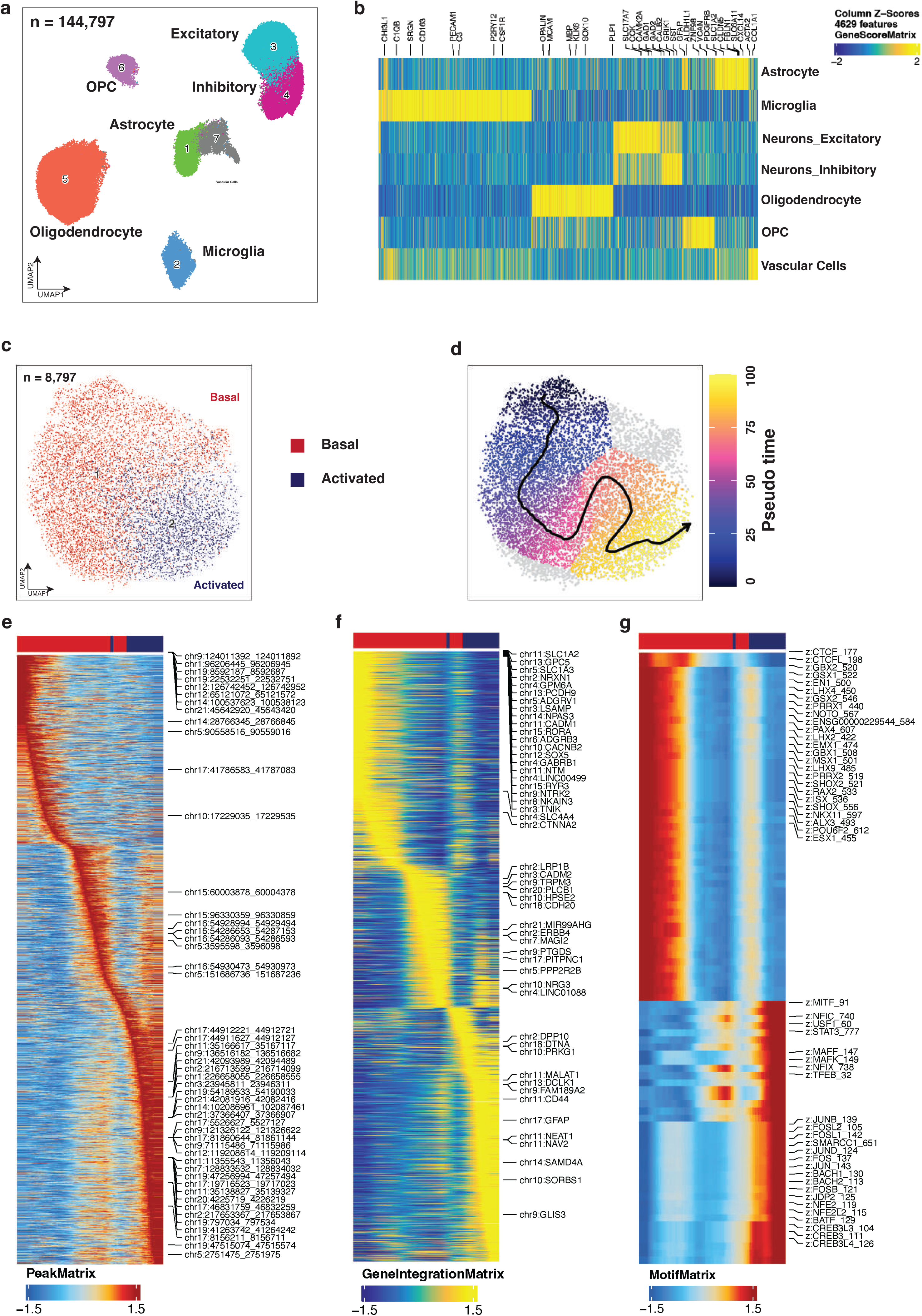
snATAC-seq recapitulates astrocyte heterogeneity and supports a basal-to-activated regulatory axis. a, UMAP embedding of the whole-brain snATAC-seq atlas after quality control, showing the major brain cell classes across filtered nuclei (n = 144,797). b, Marker gene-score heatmap used to annotate excitatory neurons, inhibitory neurons, oligodendrocyte lineage cells, astrocytes, microglia, and vascular-related cells. c-d, Astrocyte-focused snATAC-seq analysis of 8,797 nuclei showing RNA-guided label transfer of basal and activated astrocyte states (c) and arrangement along an inferred pseudotime (d). e-g, Trajectory-ordered heatmaps showing changes for accessibility of peaks correlated with the pseudotime (e), imputed or true RNA expression following snRNA and snATAC-seq integration (f), and bias-corrected, per-cell transcription factor (TF) activity scores (g).

To further dissect astrocyte-specific vulnerability, we subsetted astrocyte nuclei from the snATAC-seq dataset and re-embedded these cells after excluding samples with low astrocyte yield or quality, resulting in 8,797 astrocytes from 18 samples with well-mixed distributions across diagnosis, brain region, and individual samples (**Extended Data Fig. 5g**). Astrocyte states were annotated by integrating chromatin accessibility profiles with matched snRNA-seq references, identifying basal and activated-like states with high confidence (**Fig. 5c; Extended Data Fig. 5h**). In contrast to the six transcriptionally defined astrocyte states, chromatin accessibility profiles resolved only these two major states (**Extended Data Fig. 5h**). This difference likely reflects either limited resolution of snATAC-seq for finer state distinctions or convergence of multiple transcriptional states onto broadly similar regulatory landscapes.

Pseudotime analysis organized astrocytes along a continuous basal-to–activated-like axis, consistent with the trajectory inferred from matched snRNA-seq data (**Fig. 5d**). This gradient was evident across accessibility-based trajectory heatmaps, integrated gene activity profiles, and transcription factor (TF) deviation scores (**Fig. 5e-g**). At the level of gene activity, basal astrocytes were characterized by high *SLC1A2*, *SLC1A3*, and *ADGRV1*, whereas activated-like states showed increased *GFAP*, *DPP10*, and *DCLK1* (**Fig. 5f**). These markers were independently identified in the snRNA-seq analysis (**Fig. 1d**) and partially validated at the RNA and protein levels (**Fig. 2c-d**), supporting a coordinated transition in gene regulatory activity across the trajectory. To link transcriptional changes to regulatory mechanisms, we correlated TF gene activity with motif deviation scores across pseudotime, identifying 39 gene–motif pairs with concordant dynamics (**Extended Data Fig. 5i**). Basal states were enriched for regulators such as RORA and RORB, whereas progression along the trajectory was associated with increased activity of stress- and reactivity-associated TF programs, including AP-1, CEBP, STAT, and BACH families. Together, these results indicate that astrocyte state transitions are accompanied by progressive remodeling of the underlying regulatory landscape.

At both the single-cell and donor levels, astrocyte pseudotime distributions differed by brain region, with AMY astrocytes generally shifted toward higher pseudotime values compared to those in the PFC, consistent with the earlier and more prolonged involvement of limbic regions such as the amygdala in Down syndrome–associated Alzheimer’s disease pathology relative to neocortical regions including the PFC (**Extended Data Fig. 6a)**^2^. This regional difference was most pronounced in basal astrocytes, whereas activated-like states showed more comparable distributions across regions (**Extended Data Fig. 6b**). Differences between DS and DSAD were more subtle and context-dependent, with DSAD samples generally exhibiting a modest shift toward higher pseudotime across most regions and states, but an opposite trend observed in PFC basal astrocytes (**Extended Data Fig. 6c**). To independently validate this trajectory, we applied an external astrocyte activation gene signature^24^ and observed a highly concordant gradient across cells, with activation scores closely tracking pseudotime values (**Extended Data Fig. 6d**).

Together, these results define a region-dependent trajectory of astrocyte state progression and provide a framework for examining disease-associated changes, particularly within PFC basal astrocytes.

### Stress and Inflammation-related Transcription Factors Show Reduced Motif Accessibility in DSAD Relative to DS in PFC Basal Astrocytes

To identify cis-regulatory mechanisms underlying the transcriptional dysregulation observed in PFC basal astrocytes, we performed differential chromatin accessibility analysis between DS and DSAD. This analysis revealed widespread remodeling of the accessible chromatin landscape in DSAD, with a global shift toward increased accessibility. In total, 13,038 regions were differentially accessible, including 8,855 enriched in DSAD and 4,183 enriched in DS (**Fig. 6a; Extended Data Fig. 7a**).

**Figure 6.**
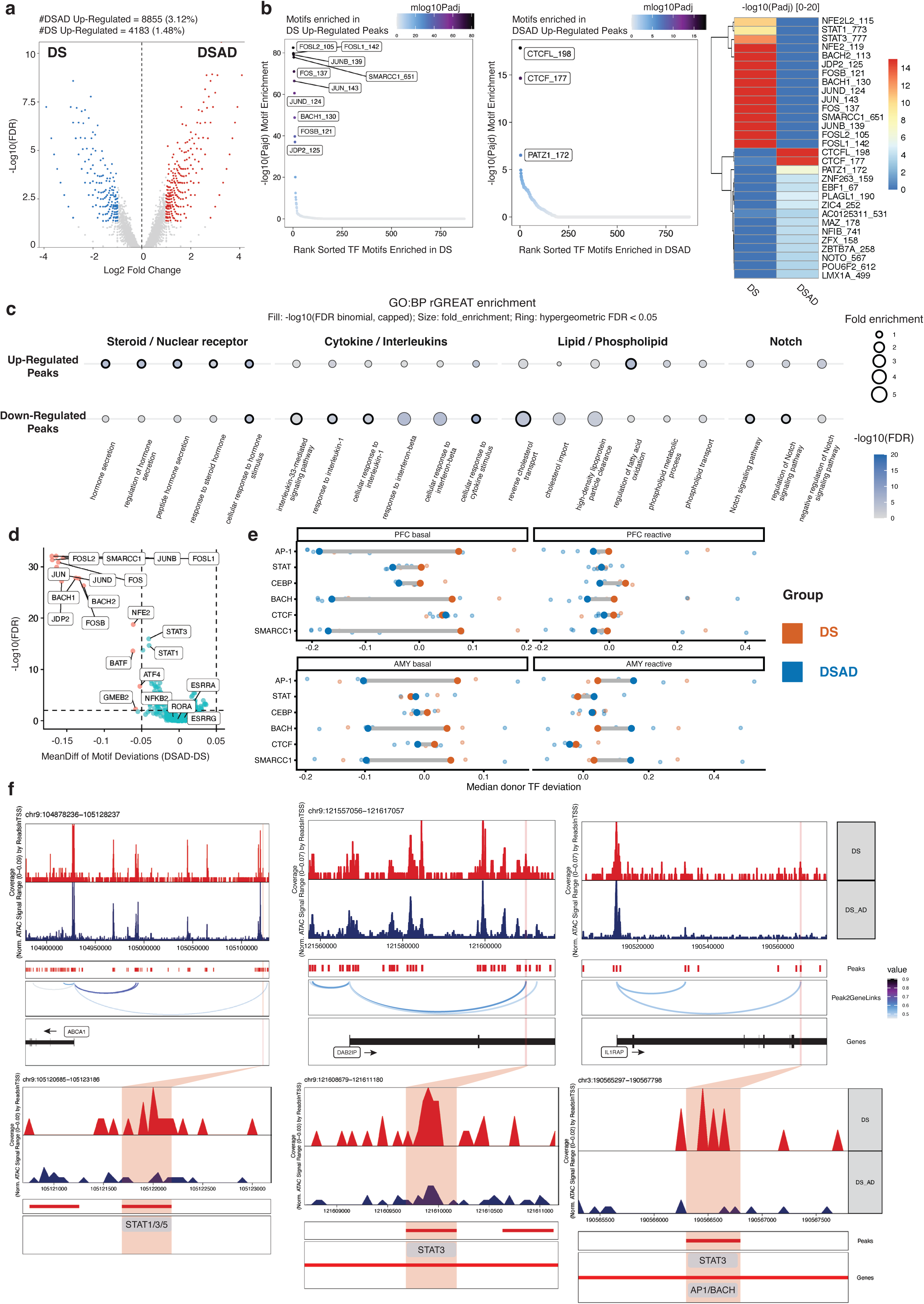
Cis-regulatory remodeling in DSAD PFC basal astrocytes. a, Differential accessibility analysis in PFC basal astrocytes identifying diagnosis-associated differentially accessible peaks (DAPs) between DS and DSAD. b, Motif enrichment analysis of DS-enriched and DSAD-enriched DAPs, including AP-1, BACH, and STAT family motifs among DS-enriched peaks and CTCF/CTCFL motifs among DSAD-enriched peaks. c, rGREAT functional enrichment analysis of diagnosis-associated DAPs based on the GO Biological Process database, with a focus on cytokine, interferon, lipid transport, cholesterol transport, Notch-related, steroid, hormone, and nuclear receptor-associated terms. d, ChromVAR-based differential motif deviation analysis in PFC basal astrocytes. TFs of interest are labeled. e, Dumbbell plot comparing median donor deviations of a selected group of TFs between DS and DSAD across the two astrocyte states and the two brain regions. f, Representative browser tracks and linked candidate cis-regulatory elements at loci including ABCA1, DAB2IP, and IL1RAP, integrating differential accessibility, motif content, and peak-to-gene linkage.

To link diagnosis-associated epigenetic changes to transcriptional alterations in PFC basal astrocytes, we examined chromatin accessibility at promoters of differentially expressed genes between DS and DSAD (**Extended Data Fig. 7b**). Consistent with the transcriptomic analysis, key genes implicated in astrocyte dysregulation, including *ABCA1*, *STAT1*, *IL1RAP*, and *ESRRG*, exhibited concordant changes in promoter accessibility and gene expression.

Motif enrichment analysis of DS- and DSAD-enriched differentially accessible regions identified distinct regulatory programs across conditions (**Fig. 6b**). DS-enriched regions were preferentially associated with stress- and inflammation-responsive motifs, including AP-1 (FOS–JUN), BACH, and STAT families. In contrast, DSAD-enriched regions showed increased enrichment for motifs linked to chromatin organization, including CTCF/CTCFL (**Fig. 6b**). These patterns suggest a relative depletion of stress- and inflammatory-response regulatory elements in DSAD, accompanied by increased accessibility at chromatin architecture–associated sites, consistent with the altered positioning of DSAD basal astrocytes along the pseudotime trajectory (**Extended Data Fig. 6c**). Notably, this difference is not attributable to subtype binarization or differential enrichment of activated-like astrocytes, as DSAD astrocytes exhibit lower activation scores than DS when considering all PFC astrocytes collectively (**Extended Data Fig. 6d**).

To place these differentially accessible peaks in biological context, we linked DS- and DSAD-enriched peaks to functional programs (**Fig. 6c; Extended Data Fig. 7c**). DS-enriched peaks were associated with cytokine and interferon signaling, including IFN–STAT-related pathways, as well as lipid, phospholipid, and cholesterol transport processes. In contrast, DSAD-enriched regions were preferentially associated with steroid/hormone and nuclear receptor-related programs (**Fig. 6c; Extended Data Fig. 7c**). Notably, Notch-related terms were enriched among DS-enriched peaks, consistent with reduced JAG1–NOTCH2 co-expression between astrocytes and vascular cells observed in the matched snRNA-seq analysis (**Fig. 6c, Extended Data Fig. 4i**). Together, these results support a shift in PFC basal astrocytes from cytokine and lipid-associated regulatory programs toward nuclear receptor and steroid-dominated programs in DSAD.

We next assessed whether these motif-level differences are reflected in TF activity at the chromatin level. Both chromVAR motif deviations and motif footprinting analyses indicated reduced activity of stress- and immune-responsive TF programs in DSAD relative to DS, including AP-1, BACH and STAT families, supporting the robustness of these findings in PFC basal astrocytes (**Fig. 6d; Extended Data Fig. 8a-b**). In contrast, these differences were weaker or less consistent in activated-like astrocytes and in the AMY (**Fig. 6e; Extended Data Fig. 8c**), indicating that this regulatory shift is not a global astrocyte feature but is most pronounced in the vulnerable PFC basal astrocyte population.

Finally, integrating differentially accessible peaks with motif annotations and peak-to-gene links prioritized candidate cis-regulatory elements (CRE) associated with differential gene expression. For example, DS-enriched peaks linked to lipid/cholesterol transport and cytokine receptor genes, including *ABCA1*, *DAB2IP* and *IL1RAP*, contained motifs for STAT, AP-1, and BACH families (**Fig. 6f**). These motif-informed peak-to-gene links suggest that distinct gene modules may be regulated by different combinations of TFs, with some loci associated primarily with STAT-family factors, such as *ABCA1* and *DAB2IP*, and others showing coordinated enrichment of STAT, AP-1 and BACH motifs, such as *IL1RAP*. Notably, these genes were also consistently identified across transcriptomic and pathway analysis, supporting their relevance to astrocyte dysregulation. Comparison of the corresponding loci across astrocyte states and regions further suggested that these accessibility differences were most pronounced in PFC basal astrocytes, whereas shifts were less evident in activated-like astrocytes and in the amygdala (**Extended Data Fig. 9a-c**).

## Discussion

By integrating matched single-nucleus RNA-seq and ATAC-seq profiles across the prefrontal cortex (PFC) and amygdala (AMY) from individuals with Down syndrome (DS) and Down syndrome-associated Alzheimer’s disease (DSAD), we identify a region- and cell state-specific pattern of vulnerability associated with disease progression. Rather than global changes across cell types or brain regions, the dominant signal converges on astrocytes in the PFC, driven by coordinated depletion and dysregulation of basal astrocyte states. Across modalities, these changes are characterized by loss of support-related transcriptional programs together with remodeling of the underlying regulatory landscape. Collectively, these findings support a model in which astrocyte dysfunction in DSAD arises not from excessive activation, but from a failure to engage appropriate stress- and inflammatory-response programs.

The preferential vulnerability of PFC basal astrocytes is notable given the central role of this region in higher-order cognitive processes, including executive function and working memory^25^. These findings raise the possibility that selective disruption of glial support states in the PFC contributes directly to the cognitive decline that distinguishes DSAD from cognitively resilient DS. More broadly, astrocytes are increasingly recognized as regionally specialized and transcriptionally tuned to support local neuronal circuits^26^, suggesting that region-specific astrocyte dysfunction represents a key axis of selective vulnerability in DSAD and related neurodegenerative disorders.

Importantly, this vulnerability is not uniformly distributed across astrocyte states. Among the six transcriptionally defined astrocyte states, depletion is most pronounced in basal states, which are enriched for homeostatic functions including neurotransmitter buffering, lipid and cholesterol metabolism, and maintenance of tissue homeostasis. Loss of these states suggests erosion of a critical glial support layer required to sustain PFC circuit stability under pathological stress. In this context, the relative preservation or expansion of non-basal astrocyte states—here transcriptionally defined as “activated-like”—may not simply reflect classical astrogliosis, but instead occur alongside depletion of homeostatic support programs, highlighting a decoupling between astrocyte activation signatures and supportive function^27^.

By contrast, the absence of a comparable depletion signature in the AMY does not necessarily indicate reduced involvement of this region. Alzheimer-related pathology in DS is known to emerge decades before clinical onset and follows a trajectory similar to autosomal dominant AD, with early involvement of limbic regions and later spread to neocortical areas^28,29^. The presence of similar regulatory alterations across regions, coupled with selective depletion in the PFC, suggests that a shared astrocyte response may have region-specific consequences, with PFC astrocytes more vulnerable to loss under disease conditions. This is consistent with the later involvement of prefrontal cortex in neurofibrillary tangle progression, whereas limbic regions such as the amygdala are affected earlier^30^.

Beyond changes in abundance, surviving basal astrocytes in the PFC exhibit coordinated transcriptional reprogramming, characterized by increased steroid- and nuclear receptor–associated programs (including *ESRRG* and *RORA*) together with reduced cytokine signaling and lipid-handling pathways (including *IL33*, *IL1RAP*, *STAT1*, *ABCA1*, and *DAB2IP*). These changes indicate a shift away from tissue-supportive and immune-responsive functions toward a functionally constrained state, consistent with broader evidence that astrocytes in neurodegenerative disease can lose key homeostatic functions while engaging context-dependent adaptive or maladaptive programs^26,31,32^.

Consistent with this model, snATAC-seq analysis reveals that these transcriptional changes are accompanied by broad cis-regulatory remodeling, including reduced accessibility of immune- and stress-responsive transcription factor programs such as AP-1, STAT, and BACH families. Integration of chromatin accessibility with peak-to-gene linkage implicates regulatory disruption at loci including *ABCA1*, *DAB2IP*, and *IL1RAP*, suggesting candidate TF–CRE–gene networks through which stress-responsive transcription factors may help maintain basal astrocyte programs. Together, these findings support a model in which astrocytes in DSAD adopt an epigenetically constrained, hypo-responsive state, limiting their ability to activate protective transcriptional programs in response to pathological stress.

Notably, astrocyte dysfunction in the PFC occurs alongside coupled depletion of endothelial cells and disruption of astrocyte–vascular signaling, including reduced JAG1–NOTCH2 interactions between astrocytes and vascular-associated cells. Given the roles of astrocytes and vascular compartments in maintaining blood–brain barrier integrity, metabolic exchange, and tissue homeostasis^26,32^, these findings raise the possibility that DSAD-related pathology involves disruption of a broader glial–vascular support niche, rather than astrocyte dysfunction in isolation.

By leveraging DS as a genetically defined model of Alzheimer-related neurodegeneration, this study provides a framework to identify regulatory mechanisms of cellular vulnerability that may be obscured in more heterogeneous sporadic AD populations. Recent single-cell studies of AD and DSAD have revealed diverse astrocyte states, including homeostatic, intermediate, reactive, and context-dependent phenotypes that vary across brain regions and disease stages^13,33^. Our findings complement this framework by highlighting a distinct mode of astrocyte dysfunction characterized by reduced responsiveness and regulatory constraint, suggesting that, in addition to reactive remodeling, failure of astrocytes to appropriately engage protective stress and inflammatory programs may represent an underappreciated mechanism contributing to neurodegenerative progression.

Several limitations should be considered when interpreting these findings. First, the number of DS donors was limited, reflecting the rarity of well-characterized tissue from individuals with DS without dementia^13,33^. Although profiling multiple brain regions increases the number of analyzed samples, donor-level statistical power remains constrained. Second, our cross-sectional design does not resolve the temporal progression of astrocyte state changes, and the observed depletion and reprogramming may represent both drivers and consequences of disease progression. Third, changes in astrocyte proportions inferred from single-nucleus data cannot distinguish between true cell loss and transitions into other states, and may also be influenced by technical factors such as nuclei recovery and tissue quality. Finally, both snRNA-seq and snATAC-seq data from postmortem tissue are subject to technical variability and sparse signal, which may limit sensitivity for detecting subtle or heterogeneous effects.

In summary, by directly comparing DS and DSAD within a shared genetic background across two brain regions, we identify a region-specific vulnerability centered on PFC basal astrocytes, characterized by coordinated depletion, transcriptional reprogramming, and cis-regulatory remodeling. These findings define a previously unrecognized astrocyte state marked by epigenetic constraint and reduced responsiveness, and provide a conceptual framework for understanding how failure of astrocyte support programs may contribute to neurodegenerative progression.

## Acknowledgements

We thank past and present members of the Kellis, Head, Tsai, and Yang laboratories for thoughtful scientific discussions. This work was supported by The Alana Down Syndrome Center at MIT (M.K., L.T., R.F.), the Alana USA Foundation, Inc, NINDS Neuroimmunology Training Program T32 NS121727-01 (C.St.), NIH/NIA U19AG068054 (ABC-DS) and NIH/NIA P30AG066519 (ADRC) to L.He., and startup funding from the School of Arts and Sciences and the Human Genetics Institute of New Jersey at Rutgers, The State University of New Jersey (J.Y.).

## Author Contributions

This study was designed and directed by J.Y., M.K. and L.He. S.W., and L.H. compiled clinical information. K.G., L.Ho, and S.W. coordinated sample acquisition. K.G. performed scRNA-seq and snATAC-seq experiments. J.Y., C.Su., R.T. and N.S. performed data processing and computational analysis. C.St. performed immunofluorescence and imaging analysis. L.Ho and A.R. designed and performed the RNAScope analysis. R.F. and L.T. provided scientific feedback. C.Su., R.F. and J.Y. wrote the manuscript.

## Materials and Methods

### Patient cohorts for single-cell analysis

The fresh-frozen human brain tissue samples were obtained from the University of California, Irvine Alzheimer’s Disease Research Center (ADRC)’s Tissue Repository under IRB #20141526. Our study focused on brain samples from individuals with Down syndrome with (DSAD) and without (DS) Alzheimer’s Disease (AD). AD and other neuropathological diagnoses were made using the standardized neuropathological evaluation established by the National Alzheimer’s Coordinating Center (NACC).

### Single-cell RNA-sequencing and preprocessing

We isolated single nuclei from fresh-frozen human brain tissue samples according to previously published protocol^4^. Briefly, we homogenized the tissue with 15 strokes in an ice-cold homogenization buffer, cleared the homogenate by passing it through a 40 micron filter, loaded it on top of a 30/40% Iodixanol (Opti-prep) gradient as a 25% Iodixanol solution, and centrifuged it at 10,000g for 5 min at 4°C. We washed the isolated nuclear pellet twice with ice-cold 0.04% BSA in PBS, resuspended it in 0.04% BSA in PBS at a concentration of 1,000 nuclei per uL, and immediately proceeded with the generation of single-nucleus RNA (snRNA) libraries using the droplet-based sequencing technology and the Chromium Single Cell 3’ RNA reagent kit v3.1 from 10X Genomics. We profiled 5,000-6,000 nuclei per sample according to the manufacturer’s protocol. The generated cDNA libraries were indexed, pooled, and sequenced in batches using the NovaSeq 6000 S2 system and reagent kits (100 cycles; Illumina). We received the sequences, mapped the reads against the GRCh38-2020-A human reference genome, and quantified the read counts using Cell Ranger pipeline version 5.0.0 (10x Genomics).

### Single-nucleus ATAC-sequencing and preprocessing

We used the remaining nuclei to perform snATAC-seq using the Chromium single cell ATAC v1 chemistry from 10x Genomics. Briefly, we transferred and resuspended the nuclei in the Diluted Nuclear buffer at a concentration that allowed us to target 5,000 nuclei per sample. Next, we generated indexed libraries following the manufacturer’s protocol, pooled them and sequenced them using the Novaseq 6000 S2 (Illumina) technology. We aligned and quantified the libraries using Cell Ranger ATAC v1.2.0 (10x Genomics) with the GRCh38-1.2.0 reference.

### Single-cell RNA-seq data analysis

#### Quality control, dimensionality reduction, clustering and major cell type annotation

Raw snRNA-seq count files were used to create a Seurat (V4) object for downstream analysis. During preprocessing, nuclei with fewer than 500 UMIs, fewer than 200 detected genes, 7,000 or more detected genes, or at least 15% mitochondrial transcripts were excluded. Mitochondrial transcript percentage was calculated for each nucleus and used together with nCount_RNA and nFeature_RNA as standard quality control metrics. After quality control, the filtered datasets were normalized and integrated across samples using Seurat canonical correlation analysis (CCA)-based anchor integration. Dimensionality reduction, graph-based clustering, and UMAP visualization were then performed on the integrated data. UMAP embeddings of the finalized atlas were examined across diagnosis, brain region, sample, sex, APOE status, age, postmortem interval, and standard RNA quality metrics. Major cell classes were annotated using canonical marker genes, and marker-gene dot plots and feature-level visualizations were used to confirm excitatory neuron, inhibitory neuron, oligodendrocyte progenitor cell, oligodendrocyte, astrocyte, microglial, and vascular-related populations.

#### Cell type-specific subclustering and subtype annotation

Astrocyte and oligodendrocyte subsets were explicitly reanalyzed by variable feature selection, PCA, shared nearest neighbor graph construction, graph-based clustering, and UMAP visualization. Cluster identities were assigned by combining marker gene expression with cluster structure, and subtype marker enrichment was assessed using Seurat FindAllMarkers function. The resulting astrocyte and oligodendrocyte subtype annotations were used in downstream analyses Neuronal subtype labels were assigned on the basis of canonical marker expression and regional transcriptional organization, using published human cortical subtype nomenclature as a reference for consensus excitatory and inhibitory neuronal identities^16,34^. Microglial subtype labels were assigned by comparing cluster marker genes and functional signatures with published human microglial state frameworks, including homeostatic, phagocytic, antigen-presentation, stress-related, DAM, brain-associated macrophage (BAM), and interferon programs^6^. The finalized neuronal and microglial subtype annotations were then used for downstream visualization, abundance analysis, and differential expression analysis..

#### Cross-dataset validation using a public euploid snRNA-seq dataset

To assess whether the major cell types and astrocyte states identified in the DS and DSAD samples were preserved in euploid brain tissue, we analyzed integrated Seurat objects containing both scDS/public euploid objects^16^ using Seurat CCA-based anchor workflow. The resulting combined object was then examined across study, brain region, sample, diagnosis, and cell type annotations to evaluate alignment of shared major cell populations across datasets. Astrocyte subtype labels were further examined in an integrated astrocyte-focused object containing both scDS and euploid astrocytes to assess whether the astrocyte states identified in the scDS cohort were reproducibly observed in the external euploid dataset.

#### InferCNV-based chromosome 21 copy-number inference

To evaluate large-scale copy-number patterns and confirm chromosome 21 trisomy, inferCNV was applied to raw count matrices from the combined scDS/euploid object. Gene ordering information was restricted to genes present in the expression matrix, and nuclei were annotated by donor-region groups. Euploid donor-region groups were used as the reference populations. inferCNV was run with ‘cutoff = 0.1’, ‘denoise = TRUÈ, and ‘HMM = TRUÈ, with both ‘cluster_by_groups = FALSÈ and ‘cluster_by_groups = TRUÈ settings.

#### CytoTRACE-based analysis of the basal-to-activated astrocyte trajectory

To examine the trajectory from basal to activated astrocyte states, we analyzed the integrated astrocyte object using CellRank. CellRank’s CytoTRACEKernel was used to compute CytoTRACE-based pseudotime (’ct_pseudotimè), and a CytoTRACE transition matrix was generated from this kernel. The resulting ‘ct_pseudotimè values were visualized on the astrocyte UMAP embedding and summarized across astrocyte states to assess progressive state transitions.

#### Cell type and subtype abundance analyses

Sample-level cell type abundances were calculated as the number of nuclei assigned to a given cell type divided by the total number of nuclei in each sample-region group. Cell subtype abundances were calculated analogously. For subtype-focused analyses, subtype proportions were additionally normalized to the abundance of the corresponding major cell class, including excitatory neurons, inhibitory neurons, astrocytes, oligodendrocytes, and microglia. Sample-level excitatory-to-inhibitory neuron ratios and OPC-to-oligodendrocyte ratios were also calculated from the corresponding cell count matrices. Diagnostic differences in cell type and subtype proportions were assessed separately within each brain region using Wilcoxon rank-sum tests. For comparisons involving public euploid samples, sample-level cell type abundances and inhibitory-to-excitatory neuron ratios were also calculated on the combined scDS/euploid object and compared between studies within each brain region.

#### Cell type and subtype correlation analyses

To examine cellular composition across individuals, sample-level matrices of cell type and cell subtype abundances were compared separately for AMY and PFC samples. Correlation analyses were performed on both raw abundance matrices and proportion-normalized matrices. Pearson correlation coefficients were computed for all pairs of cell types or subtypes across samples, significance was evaluated using ‘cor.test’, and correlation matrices were visualized after clustering of the correlation structure.

#### Differential expression analyses across five statistical models

To identify transcriptional differences between DS and DSAD samples while accounting for the limited DS sample size, differential expression analyses were performed separately within each brain region and cell subtypes using five statistical models. Four models were fit using NEBULA on raw count matrices with subject ID as the grouping variable and nCount_RNA as an offset, using the following design formulas: Age + Sex + PMI + DX, Sex + PMI + DX, Age + Sex + PMI + APOE + DX, and Age + PMI + APOE + DX. In parallel, a fifth model was performed using MAST as implemented in Seurat’s FindMarkers function without additional covariates. For NEBULA analyses, P values corresponding to the diagnosis term were adjusted using the Benjamini–Hochberg method. Genes with adjusted P values below 0.05 were considered significant, and effect direction was assigned according to the sign of the diagnosis-associated log fold change.

#### Pathway enrichment analyses based on consensus DEGs

To define robust diagnosis-associated transcriptional changes, genes were classified as consensus differentially expressed genes (DEGs) when they were significant across all five statistical models and showed the same direction of effect within a given brain region and cell subtype. Consensus DEG lists were generated separately for upregulated and downregulated genes in each region-subtype group. These gene lists were then submitted to Metascape^35^ for pathway enrichment analysis and network visualization.

#### Within-region ligand-receptor analysis using CellPhoneDB

To identify diagnosis-associated ligand-receptor interactions, sample-level metadata and raw gene-count matrices were generated from the snRNA-seq count matrix for matched individuals with sufficient representation of both AMY and PFC nuclei. Only matched individuals with more than 50 nuclei in each region were retained for CellPhoneDB analyses. CellPhoneDB statistical_analysis was run separately for each individual using gene symbols as input and a threshold of 0.2. Significant interactions were summarized separately for within-AMY and within-PFC comparisons. For differential interaction analyses between DS and DSAD, sample-specific interaction rank and value columns were merged across individuals, diagnosis-level mean interaction values and log fold changes were calculated, and Wilcoxon rank-sum tests were used to compare interaction ranks between diagnosis groups. Significant differential interactions were subsequently converted to directed source-target interaction tables for network visualization.

### Single-nucleus ATAC-seq data analysis

#### Quality control, dimensionality reduction, batch correction, and clustering

For whole-brain analyses: The ArchR package was used for performing our single-nucleus ATAC-seq data analysis. For the whole-brain snATAC-seq atlas, nuclei with more than 1,000 peak-region fragments and a transcription start site (TSS) enrichment score greater than 4 were retained. Doublets were then inferred using ArchR’s UMAP-based approach with 10 nearest neighbors and were filtered using a DoubletEnrichment cutoff of 1 and a filter ratio of 1. For dimensionality reduction, we applied ArchR’s iterative latent semantic indexing (LSI) to perform term frequency inverse document frequency (TF-IDF) normalization and singular value decomposition of the peak-by-cell matrix. We then extracted UMAP embeddings to visualize the dataset, then applied DBScan to obtain the final embeddings. Using marker genes from previous brain studies^5,36^, we assigned cell type labels to each cell. For astrocyte-focused analyses: After annotation, nuclei annotated as astrocytes were extracted from the whole-brain atlas and reprocessed as an astrocyte-specific ArchR project. Sample-level quality metrics were summarized for each sample-region group, including cell number, total fragments, median ReadsInTSS, median TSSEnrichment, median nucleosome ratio, and median blacklist ratio. Six low-quality sample-region groups (416_PFC, 2906_PFC, 3304_AMY, 1302_AMY, 2906_AMY, and 3110_AMY) were excluded based on low cell recovery and/or low fragment and TSS-related quality metrics. Similarly, the resulting astrocyte object was then subjected to iterative LSI on the TileMatrix followed by Harmony batch correction using patient ID as the grouping variable. UMAP embeddings were generated using 30 neighbors, a minimum distance of 0.5, and cosine distance. Clustering was performed on the Harmony-corrected embedding using the Seurat method implemented in ArchR across multiple resolutions, and the clustering solution at resolution 0.8 was selected for downstream analyses based on cluster reproducibility, adequate cell coverage, and minimal bias by patient, diagnosis, or brain region.

#### Annotation of cell types, and subtypes

For the annotation of major cell types in the whole-brain snATAC-seq atlas, we examined gene activity scores of canonical marker genes using ArchR and assigned cell type labels based on previously established brain marker sets. For astrocyte-focused analyses, astrocyte subtype annotations were transferred from the matched snRNA object to the snATAC object using ArchR’s addGeneIntegrationMatrix function, which aligns each snATAC nucleus to its most similar snRNA cell and generates a GeneIntegrationMatrix representing the predicted gene expression profile for each nucleus, based on imputation weights, the GeneScoreMatrix, and Harmony-corrected embeddings. In the unconstrained transfer, low-resolution astrocyte labels were transferred, followed by a constrained transfer within the basal and reactive subtypes by defining a groupList based on the unconstrained labels and using the higher-resolution RNA annotation. To evaluate annotation robustness, we examined the distributions of prediction scores across diagnosis and brain region, summarized both cell and donor-level prediction confidence. Because low-resolution astrocyte assignments showed higher confidence and more stable separation than the higher-resolution labels, downstream astrocyte analyses primarily used the unconstrained low-resolution groups, corresponding to basal and reactive astrocyte states.

#### Peak calling and peak matrix construction

To define accessible chromatin peaks, we utilized ArchR functions addGroupCoverages and addReproduciblePeakSet to call peaks from the astrocyte-specific ArchR project using MACS2 (v2.2.6). Pseudo-bulk coverage tracks were generated according to the Harmony-based clustering solution at resolution 0.8. Given the size of the astrocyte snATAC-seq dataset, we set maxCells = 1000 and maxReplicates = 24 in addGroupCoverages to avoid excessive downsampling while retaining all available sample contributions. Reproducible peaks were then identified across clusters using addReproduciblePeakSet with ArchR default settings. A union astrocyte peak set was subsequently used to construct the PeakMatrix with addPeakMatrix for downstream chromatin accessibility analyses. Peaks in the union set were further annotated by genomic context, including promoter, distal, intronic, and exonic categories, as well as by nearest gene and distance to the nearest transcription start site. The PFC basal astrocyte subset used in downstream analyses inherited this union peak set and PeakMatrix.

#### Cell-type proportion differences among brain regions and diagnosis

To evaluate astrocyte abundance in the context of the whole-brain snATAC-seq atlas, transferred astrocyte labels from the astrocyte-specific ArchR project were mapped back to the corresponding nuclei in the whole-brain metadata. For each patient-region sample, frequencies were calculated as the number of nuclei assigned to a given cell type or astrocyte state divided by the total number of nuclei in that sample-region. For astrocyte-state analyses, comparisons were restricted to the sample-region groups retained in the astrocyte-specific QC-filtered object. Diagnostic differences were visualized using boxplots with sample points and assessed within each brain region using Wilcoxon rank-sum tests.

#### Cell state trajectory, and activation score analyses

Like our snRNA analyses, we modeled the continuous transition from basal to reactive astrocytes, by defining an ArchR trajectory from basal to reactive on the Harmony-corrected astrocyte embedding using addTrajectory function. Trajectory profiles were then extracted from the MotifMatrix, GeneScoreMatrix, GeneIntegrationMatrix, and PeakMatrix using getTrajectory function, and trajectory heatmaps were generated to visualize changes in chromatin accessibility, gene activity, integrated gene expression, and motif deviations along the trajectory. Correlations between gene-activity and motif-deviation trajectories were computed using correlateTrajectories function.

To independently assess astrocyte activation, we utilized an activation gene set from prior studies^24^, including *GFAP*, *FABP7*, *MAOB*, *CRYAB*, *HSPB1*, *CHI3L1*, *SERPINA3*, *C3*, *TSPO*, *LCN2*, *THBS1*, *VIM*, *SOX9*, *ALDOC*, and *NES*. We applied ArchR’s function addMoudleScore to calculate the module score on the GeneIntegrationMatrix. Scores were further examined at both the cell level and donor level, with donor-level summaries calculated as median values within each patient, diagnosis, brain region, and low-resolution astrocyte state.

#### PFC basal astrocyte subset for diagnosis-focused chromatin analysis

For diagnosis-focused chromatin analyses, we subsetted low-resolution basal astrocytes from the QC-filtered astrocyte ArchR project and further restricted these nuclei to the PFC region. This PFC basal astrocyte ArchR project was used for the downstream diagnosis-associated differential accessibility, motif enrichment, TF deviation, promoter accessibility concordance, pathway enrichment, peak-to-gene linkage, and browser-track analysis.

#### Differential accessibility, motif enrichment, TF deviation, footprinting analysis

Differential analyses were performed within the PFC basal astrocyte population. Diagnosis-associated differentially accessible peaks (DAPs) between DS and DSAD were identified from the PeakMatrix using ArchR’s getMarkerFeatures function with a binomial test (groupBy = “DX”, normBy = “none”, bias = c(”TSSEnrichment”, “nFrags”), k = 100, closest = TRUE, bufferRatio = 0.8, binarize = TRUE, and maxCells = 500). DAPs were defined using a cutoff of FDR <= 0.05 and Log2FC >= 1.0. Peak annotations, including genomic category, nearest gene, and distance to the nearest transcription start site, were obtained from the ArchR union peak set.

For motif enrichment, TF motif annotations from the CIS-BP database were added to the PFC basal astrocyte ArchR project using addMotifAnnotations function, and enrichment of TF motifs in diagnosis-associated DAPs was assessed with peakAnnoEnrichment function using the same DAP cutoff. To quantify TF activity at single-cell resolution, background peaks were added with addBgdPeaks function and motif deviation scores were computed using ChromVAR as implemented in ArchR through addDeviationsMatrix function. Differential motif deviations between DS and DSAD were then assessed on the MotifMatrix using getMarkerFeatures function with a Wilcoxon test and bias correction for TSSEnrichment and log10(nFrags) (k = 100).

To summarize the chromatin deviations of selected TF programs highlighted in the trajectory and differential accessibility analyses, including AP-1, BACH, and STAT-family programs, family-level TF activity scores were derived from the QC-filtered astrocyte MotifMatrix. For each TF family of interest, motif deviation features corresponding to individual TFs within that family were identified and averaged at the cell level to generate a single family-level activity score per nucleus. These family-level scores were compared at the cell level across diagnosis, brain region, and low-resolution astrocyte state, and were additionally summarized at the donor level as median values within each patient, diagnosis, brain region, and low-resolution astrocyte state.

For footprinting analyses, diagnosis-level group coverages were generated using addGroupCoverages function (groupBy = “DX”, maxCells = 1000, maxReplicates = 10). Motif positions were identified with getPositions function, and TF footprints were computed for selected motifs representing inflammatory/stress-responsive and nuclear receptor-associated programs, including *STAT1*, *STAT3*, *NFE2L2*, *ESRRG*, *ESRRA*, *RORA*, *JUNB*, *JUND*, *FOS*, *FOSB*, *FOSL1*, *FOSL2*, *BACH1*, and *BACH2*. Footprint profiles were visualized using plotFootprints function with subtractive bias normalization (normMethod = “Subtract”), addDOC = FALSE, and smoothWindow = 5.

#### Promoter accessibility concordance with DEGs

To directly compare chromatin accessibility changes with matched transcriptional changes in PFC basal astrocytes, promoter-annotated peaks were extracted from the astrocyte union peak set using ArchR peak annotations and linked to genes based on nearest-gene assignments. Consensus DEGs from the five matched snRNA-seq models, restricted to PFC basal astrocytes, were then matched to diagnosis-associated promoter peaks assigned to the same genes. When multiple promoter peaks were assigned to a gene, promoter accessibility effects were summarized to the gene level by averaging the diagnosis-associated Log2FC values across promoter peaks. Gene-level promoter accessibility direction was then compared with the corresponding RNA differential-expression direction to classify genes as same-direction up, same-direction down, discordant, or showing no promoter accessibility change.

#### Pathway enrichment, peak-to-gene linkage, and integration with DEGs

Diagnosis-associated DAPs identified in PFC basal astrocytes were utilized to region-based functional enrichment using rGREAT on hg38 with the GO Biological Process (”GO: BP”) database. For regulatory-circle establishment, motif-containing DAPs were intersected with motif matches from the PFC basal astrocyte ArchR project and annotated to genes using rGREAT-derived region-gene associations obtained with getRegionGeneAssociations functions. DEGs from five matched snRNA-seq models (age_sex_pmi_dx, sex_pmi_dx, age_sex_pmi_apoe_dx, age_pmi_apoe_dx, and MAST) were imported and restricted to PFC basal astrocytes. Genes were summarized across models by the number of supporting models, median RNA log fold change, and concordance of direction across models. Motif-DAP-gene summaries were ranked by direction-matched DEG support, number of supporting DEG models, and distance to the transcription start site. This rGREAT-based framework was used for the initial regulatory-circle validation of STAT1/STAT3-associated DAP modules. For locus-level visualization, peak-to-gene links were subsequently computed with ArchR’s addPeak2GeneLinks function on the QC-filtered astrocyte ArchR project using the Harmony corrected reduced dimensions and retrieved with getPeak2GeneLinks function using corCutOff = 0.5 and FDR = 1e-4. These links were used to support linkage display in browser-track visualizations. Final track plots were generated in the PFC basal astrocyte ArchR project using plotBrowserTrack function grouped by diagnosis, with gene-centered windows extending 50 kb upstream and downstream and peak-centered zoomed views extending 1 kb beyond highlighted DAPs. Tracks displayed bulk accessibility, feature, loop, and gene annotations, with tileSize = 50, maxCells = 5000, minCells = 25, and log2Norm = FALSE. For comparison of prioritized candidate loci across other astrocyte populations, browser tracks were also generated for PFC reactive, AMY basal, and AMY reactive astrocyte subsets using the same candidate loci and matched display settings, together with peak-to-gene links where available.

### RNAscope

The RNAscope LS Fluorescent Multiplex Kit (cat. no. 322800) (Advanced Cell Diagnostics, Newark, CA) was used with standard pretreatment conditions. FFPE human brain tissue samples were incubated with Leica BOND Epitope Retrieval Solution 2 (ER2) at 95°C for 15 minutes. RNAscope 2.5 LS Protease III was used for 15 minutes at 40°C. Pretreatment conditions were optimized for each sample and stainings were completed using probes specific to the genes *ALDH1L1*, *GFAP*, *APOE*, *SLC1A2*, *ROBO2*, *SERPINA3*, *KCND2* and *DPP10*. Negative control background staining was evaluated using a probe specific to the bacterial *dapB* gene. Fluorescent images were acquired using a 3D Histech Panoramic Scan Digital Slide Scanner microscope using a 40x objective.

### Immunofluorescence

Immunofluorescence (IF) was conducted using formalin-fixed post-mortem human brain tissue sections from occipital and frontal cortices. All tissue was sliced at 30µm thick using the Leica Vibratome (Model# VT10005) and were stained using the following free-floating method, as described here. Tissue was washed in 1x TBS (Bioland Scientific LLC Cat# TBS01-03) for 5 minutes at RT twice. Tissue was then incubated in citrate buffer (0.0147 g/ml sodium citrate; pH 6.0; Fisher Chemical Cat# S279-500) for 30 minutes in a water bath at 80°C then allowed to cool for 10 minutes at RT. Tissue was washed in 1x TBS twice at RT for 5 minutes per wash, washed in TBS-A (1x TBS with 0.1% Triton-X100 (Fisher BioReagents Cat# BP151-100)) for 15 minutes, then incubated in TBS-B (1x TBS with 5% BSA (Gemini Bio Cat#700-100P) for 30 minutes at RT. Tissue was blocked in TBS-B with 5% normal horse serum (Vector Laboratories Cat#S-2000-20) for 1 hour at RT and then incubated in primary antibodies overnight at 4°C on an orbital shaker. Primary antibodies utilized included rabbit anti-COX3 at a 1:300 dilution (NovusBio, Cat# NBP2-97594), rabbit anti-GRID2 at a 1:200 dilution (Invitrogen, Cat# PA5-115325), rabbit anti-SULF1 at a 1:300 dilution (Invitrogen, Cat# PA5-113112), rabbit anti-DCLK1 at a 1:1000 dilution (Invitrogen, Cat# PA5-64960), rabbit anti-GPR98 at a 1:1000 (Invitrogen, Cat# PA5-84761), rabbit anti-AQP4 at a 1:1000 dilution (Invitrogen, Cat# PA5-53234), and mouse anti-GFAP at a 1:2000 dilution (Abcam, Cat# ab4648). Tissue was washed twice in TBS-A for 5 minutes per wash and then washed in TBS-B for 15 minutes. Secondary antibodies used were donkey anti-rabbit 555 (Invitrogen, Cat# A31572) and donkey anti-mouse 647 (Invitrogen, Cat# A31571) and were diluted in blocking buffer (1x TBS-B with 5% normal horse serum) at a 1:1000 dilution. Tissue was washed in 1x TBS for 5 minutes then incubated in 1x TrueBlack Lipofuscin Autofluorescence Quencher (diluted in 70% ethanol; Biotium, Cat# 23007) for 4 minutes. Tissue was washed in 1x TBS then mounted to Fisher Brand Super Frost Plus slides (Fisher Scientific Cat#12-550-15). Slides were coated with Fluoromount mounting medium with DAPI (Vector Laboratories, VECTASHIELD Vibrance antifade mounting medium Cat# H-1800) and were allowed to dry overnight in the dark before imaging.

Immunofluorescence analyses performed to assess COX3, GRID2, SULF1, DCLK1, and AQP4 localization in relation to GFAP and vasculature, identified by elongated, consistent clusters of elliptical nuclei stained with DAPI, were conducted using one representative case per diagnostic group, including DS and DS with AD groups, following imaging with a Zeiss LSM 900 with Airyscan confocal microscope at 20x objective.

### Imaging and 3D Rendering

Slides were imaged using a Zeiss LSM 900 with Airyscan confocal microscope at 20x magnification, capturing Z-stack images with 1 µm step intervals across a total depth of 10 µm within 30 µm thick tissue slices to minimize background. The Airyscan processing feature was applied to each 1 µm slice to enhance resolution. Prior to imaging, laser power and master gain settings were optimized for each channel: DAPI was set to 3% laser power and 850 V gain with a wavelength range of 400-459 nm; GFP (for GFAP) was set to 1.6% laser power and 800 V gain with a wavelength range of 485-548 nm; and Cy5 (for all other markers) was set to 3% laser power and 850 V gain with a wavelength range of 626-700 nm. Laser wavelength parameters were adjusted to minimize spectral overlap and improve signal specificity. Post-imaging, white light laser exposure was fine-tuned for each channel in the frontal and occipital cortices, with specific settings for each channel. Finally, the orthogonal array feature was applied to compress the Z-stack images.

Confocal z-stack images processed using IMARIS 10.1.0 software. For 3D rendering, the ‘Surface Rendering’ feature was applied to each channel individually. For example, the DAPI channel was rendered by selecting the ‘Surface’ feature and adjusting the surface representation incrementally to ensure accuracy. The channel color was set to blue, and the final rendering was saved. This procedure was consistently applied across all channels to maintain uniformity in 3D rendering across diagnostic groups.

## Data availability

The original data are available at GEO (GSEXXXXXX). The processed data are available on Zenodo (URL).

## Code availability

All codes that are necessary to reproduce all the results in the paper are implemented in Python and R and are publicly available at GitHub (https://github.com/YangLabRutgers/scDS).

## Supplemental information

**Supplementary Table 1. Marker genes for major cell types and annotated cell subtypes in the snRNA-seq atlas.** See Fig. 1 and Extended Data Fig. 1.

**Supplementary Table 2. Marker genes for the six astrocyte states identified in the snRNA-seq dataset.** See Fig. 2 and Extended Data Fig. 2.

**Supplementary Table 3. Sample-level cell type and cell subtype abundances in the snRNA-seq dataset.** See Fig. 3 and Extended Data Fig. 3.

**Supplementary Table 4. Differentially expressed genes between DS and DSAD across five statistical models, brain regions, and cell subtypes.** See Fig. 4 and Extended Data Fig. 4.

**Supplementary Table 5. Sample metadata and QC metrics for whole-brain and astrocyte-focused snATAC-seq analyses.** See Fig. 5 and Extended Data Fig. 5.

**Supplementary Table 6. Sample-level cell type and astrocyte-state abundances in the snATAC-seq dataset**. **See Extended Data Fig. 5.**

**Supplementary Table 7. Diagnosis-associated differentially accessible peaks in PFC basal astrocytes.** See Fig. 6 and Extended Data Fig. 7.

**Supplementary Table 8. Motif enrichment and differential ChromVAR transcription factor activity in PFC basal astrocytes.** See Fig. 6, and Extended Data Fig. 8.

**Supplementary Table 9. Functional enrichment of diagnosis-associated accessible peaks in PFC basal astrocytes.** See Fig. 6, and Extended Data Fig. 7.

**Supplementary Table 10. Integrated motif-peak-gene regulatory links in PFC basal astrocytes**. See Fig. 6.

**Supplementary Table 11. Donor-level pseudotime and external activation scores across brain regions, diagnoses, and astrocyte states in the snATAC-seq dataset.** See Extended Data Fig. 6 and Extended Data Fig. 8.

---

**Extended Data Fig. 1.**
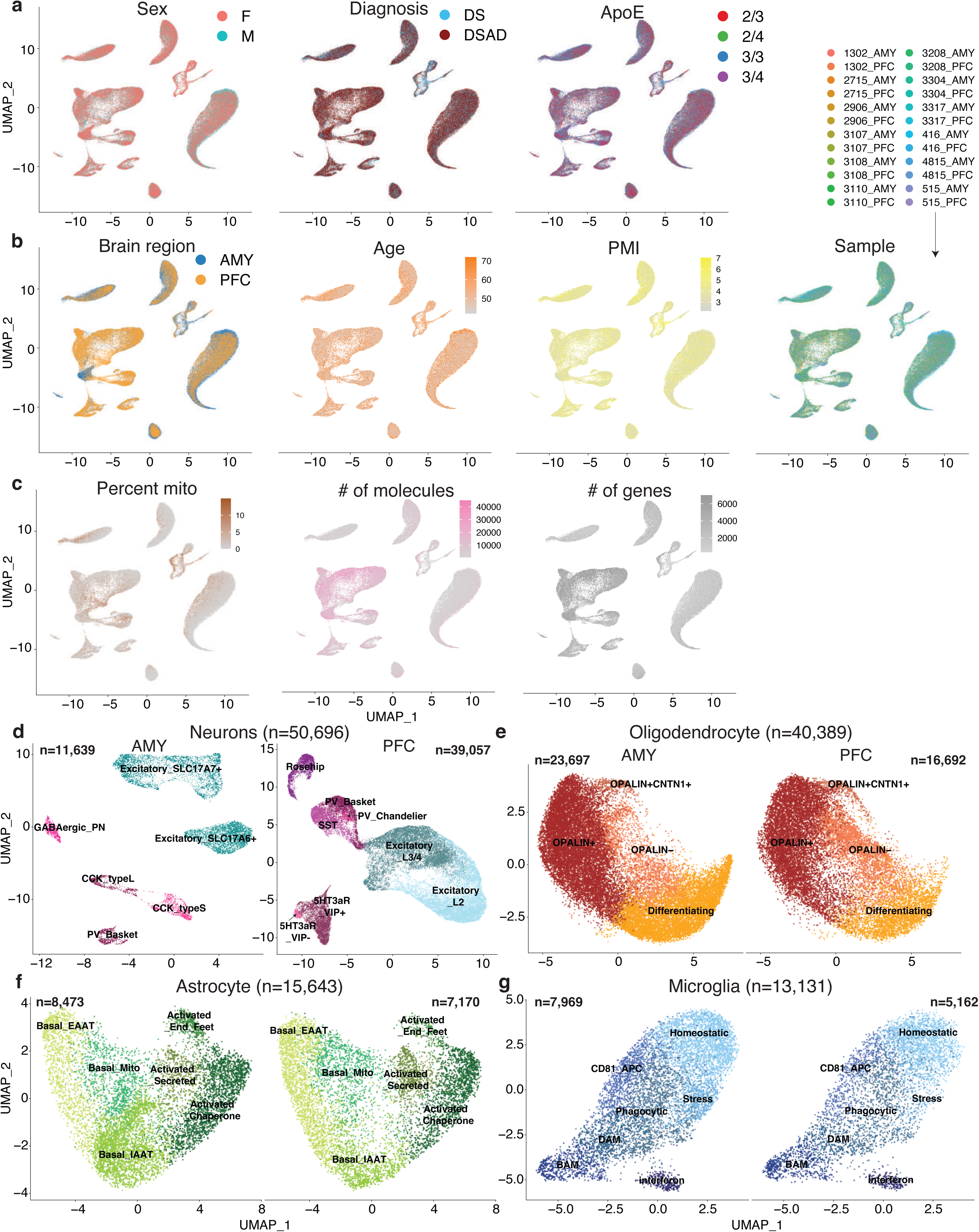
Covariates and sub-clustering across the snRNA-seq atlas. a-c, UMAP embeddings of the integrated snRNA-seq atlas, showing nuclei from the prefrontal cortex (PFC) and amygdala (AMY) colored by sample covariates and quality-control variables. a, Sex, diagnosis, and APOE status. b, Brain region, age, post-mortem interval (PMI), and sample identity. c, Mitochondrial read fraction, number of molecules, and number of genes detected per nucleus. d-g, UMAP embeddings of sub-clustered major cell classes shown separately by brain region. d, Excitatory and inhibitory neuronal subtypes. e, Oligodendrocyte subtypes. f, Astrocyte states. g, Microglial states. Labels indicate the subtype or state assignments used in downstream analyses.

**Extended Data Fig. 2.**
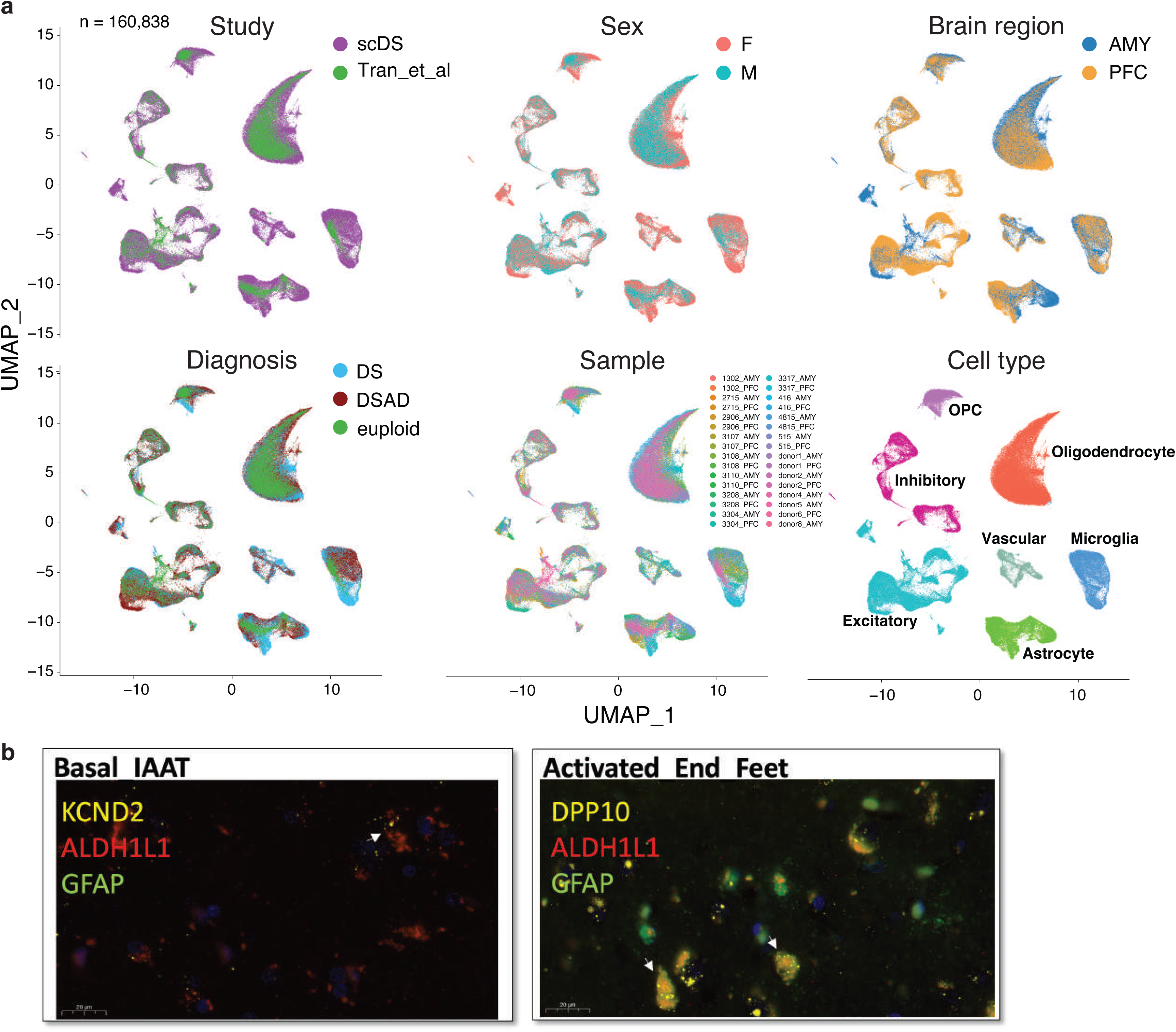
Integration of trisomic and euploid snRNA-seq datasets and additional RNAscope images. a, UMAP embeddings of the integrated dataset combining the trisomic cohort with the external euploid reference dataset, colored by study, sex, brain region, diagnosis, sample, and cell type. The same joint embedding is shown repeatedly to allow comparison across annotations. b, Representative RNAscope images for additional astrocyte-state markers not shown in Fig. 2c. Left, KCND2 with ALDH1L1 and GFAP for basal IAAT astrocytes. Right, DPP10 with ALDH1L1 and GFAP for activated end-feet astrocytes. Signal colors correspond to the probe labels shown in each panel. Scale bars, as indicated in the figure.

**Extended Data Fig. 3.**
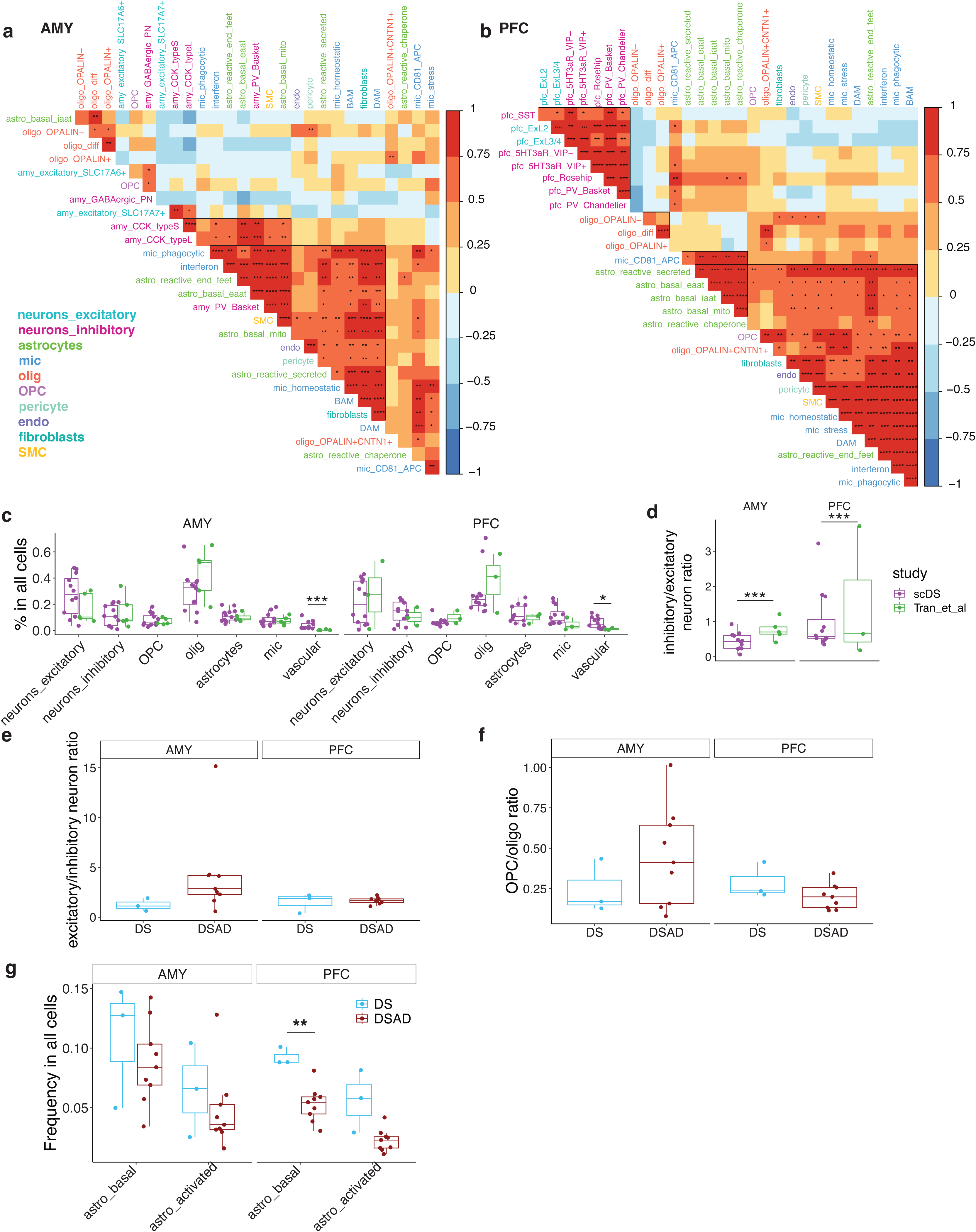
Additional cell type and subtype proportion differences between DS and DSAD. a,b, Heatmaps of donor-level correlations in subtype proportions within AMY (a) and PFC (b). Each cell shows the correlation coefficient between two subtypes across donors. c, Donor-level cell-type proportions in the trisomic cohort and the euploid reference cohort, shown separately by brain region. d, Donor-level inhibitory-to excitatory neuron ratios in the trisomic cohort and the euploid reference cohort, shown separately by brain region. e,f, Donor-level excitatory-to-inhibitory neuron ratios (e) and OPC-to-oligodendrocyte ratios (f) comparing DS and DSAD within each brain region. g, Donor-level astrocyte-state proportions comparing DS and DSAD within AMY and PFC. Boxplots summarize donor-level distributions, and points denote individual donors. Statistical comparisons are indicated where shown.

**Extended Data Fig. 4.**
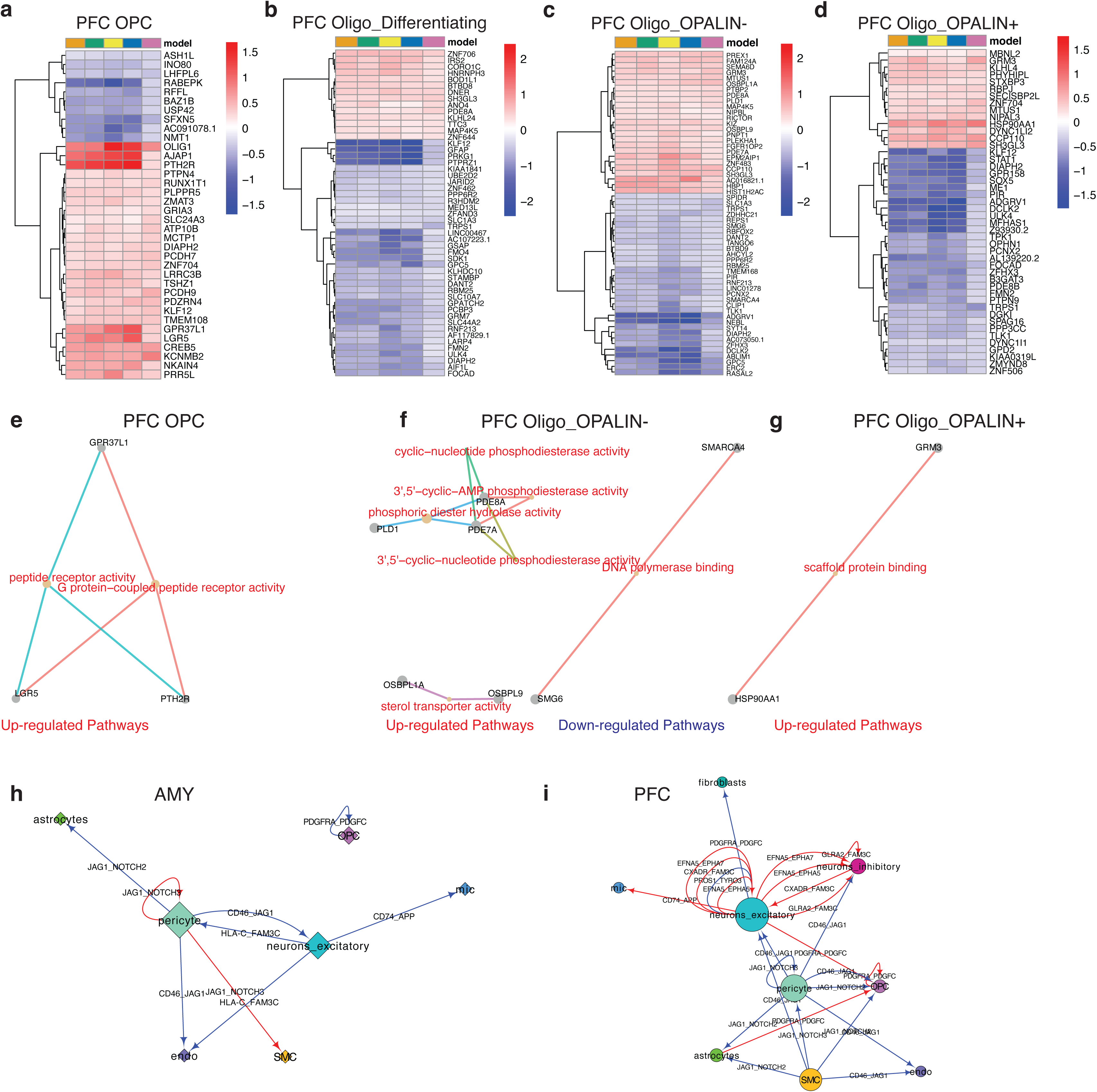
Additional DEGs and cell-cell communication results comparing DS and DSAD. a-d, Heatmaps of consensus differentially expressed genes (DEGs) for additional cell populations, shown as DSAD relative to DS. a, PFC OPC. b, PFC differentiating oligodendrocytes. c, PFC OPALIN- oligodendrocytes. d, PFC OPALIN+ oligodendrocytes. e-g, Network summaries of enriched gene-set terms for PFC OPC (e), PFC OPALIN- oligodendrocytes (f), and PFC OPALIN+ oligodendrocytes (g). h,i, CellPhoneDB-based within-region cell-cell communication networks in AMY (h) and PFC (i). Nodes represent cell types, and edges represent ligand-receptor interactions detected in each region, with edge colors shown as indicated in the figure.

**Extended Data Fig. 5.**
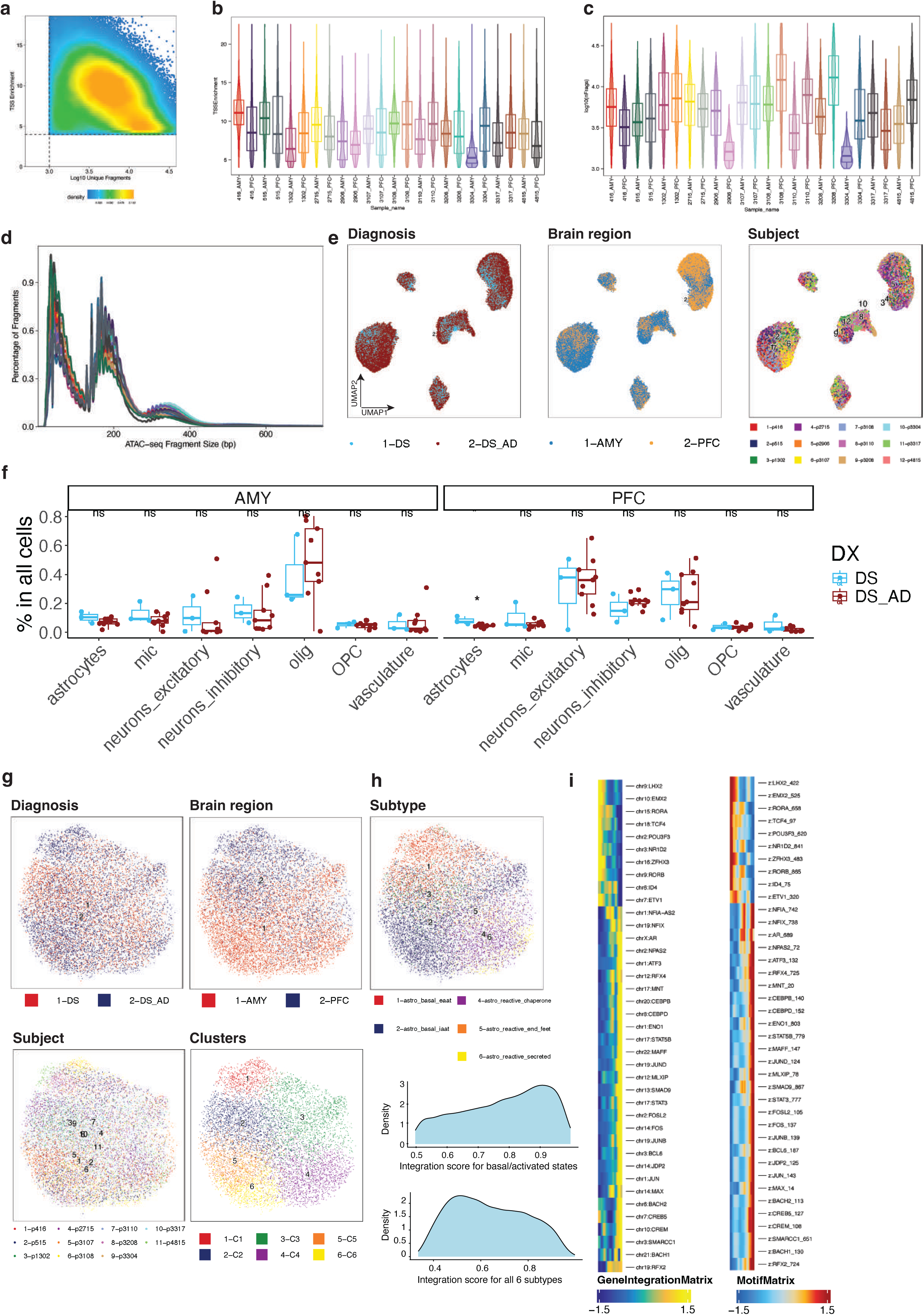
Additional snATAC-seq quality-control, integration, and astrocyte-state analyses. a, Scatter plot of transcription start site (TSS) enrichment versus unique fragment count for nuclei in the wholebrain snATAC-seq dataset, with the quality-control thresholds used for filtering indicated by dashed lines. b,c, Sample-level distributions of TSS enrichment (b) and unique fragment counts (c) after filtering. d, Fragmentsize profiles across samples, showing nucleosomal periodicity. e, UMAP embeddings of the whole-brain snATAC-seq atlas colored by diagnosis, brain region, and donor. f, Donor-level cell-type proportions in the whole-brain snATAC-seq atlas, comparing DS and DSAD within AMY and PFC. g-h, Astrocyte-focused snATAC-seq UMAP embeddings colored by diagnosis, brain region, donor, unsupervised clustering and transferred subtype label at high resolution; density plots of prediction confidence for both low and high resolutions are shown below. i, Heatmaps from the correlated GeneIntegrationMatrix and MotifMatrix arranged along the astrocyte trajectory.

**Extended Data Fig. 6.**
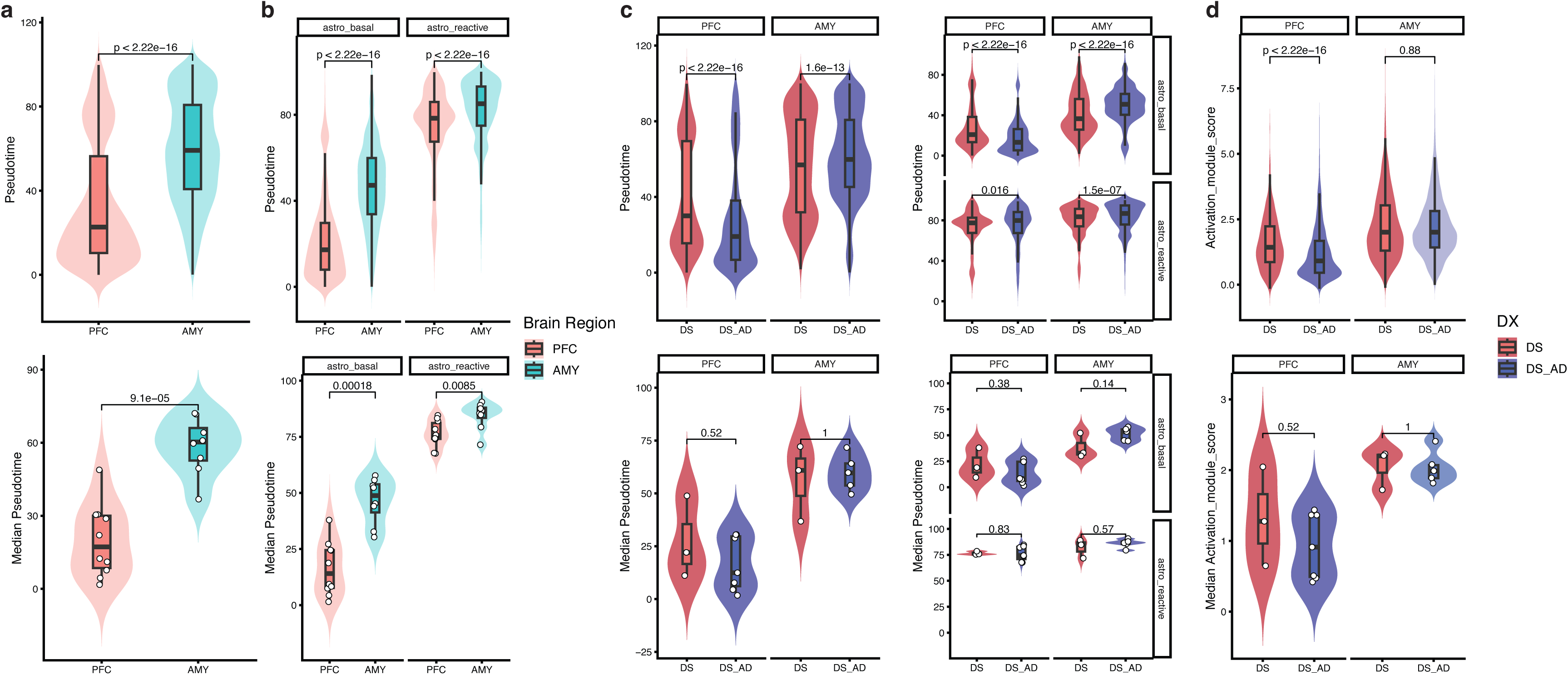
Additional astrocyte pseudotime, external activation-score, and transcription factor activity analyses. a, Cell-level and donor-level pseudotime comparisons between PFC and AMY across astrocytes. b, Cell-level and donor-level pseudotime comparisons between PFC and AMY shown separately for basal and reactive astrocytes. c, Cell-level and donor-level pseudotime comparisons between DS and DSAD, shown separately for PFC and AMY. d, External astrocyte activation-signature scores at the cell level and donor level, shown separately for PFC and AMY with comparisons between DS and DSAD.

**Extended Data Fig. 7.**
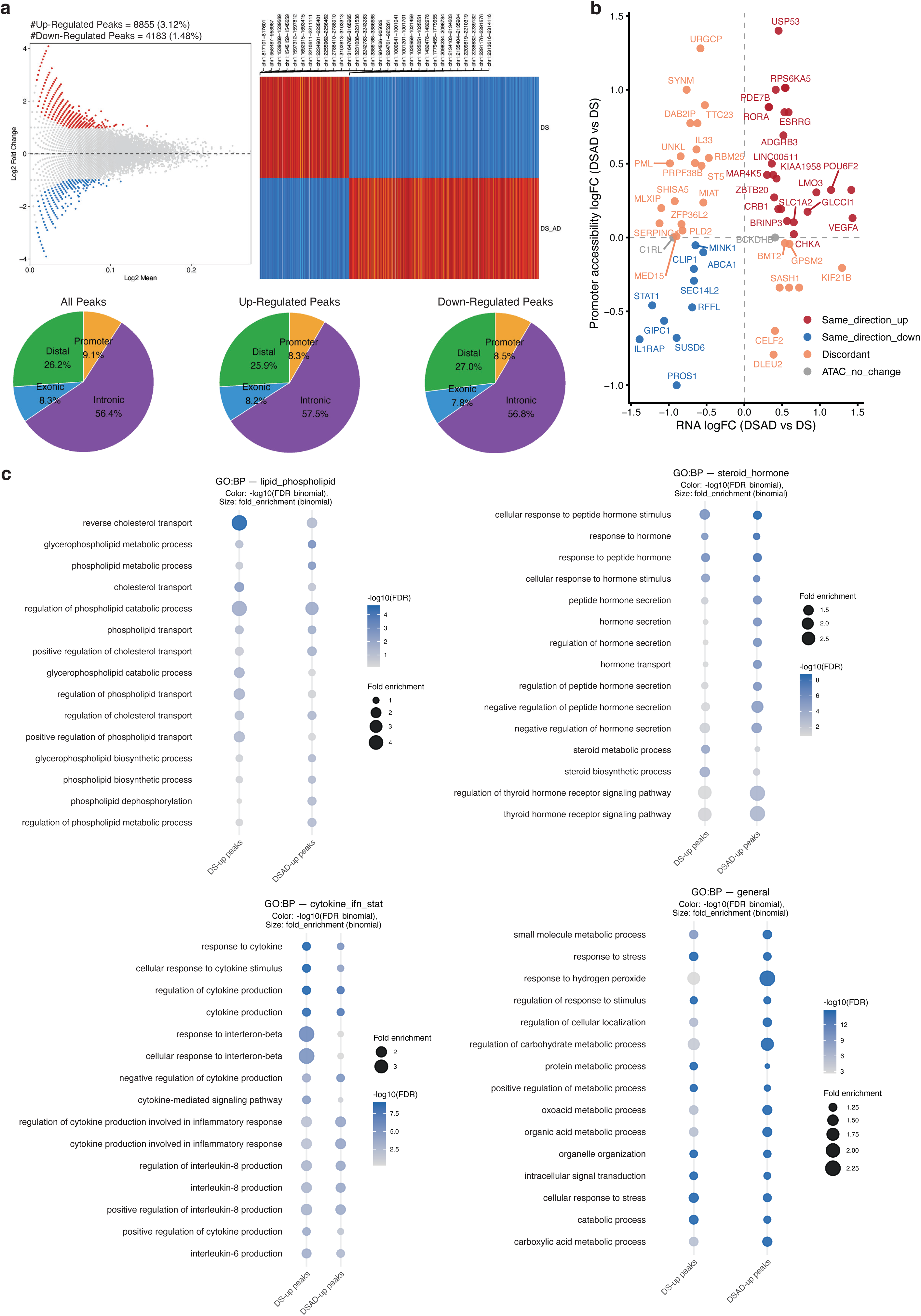
Peak-level summaries, promoter accessibility concordance, and functional annotation of diagnosis-associated DAPs in PFC basal astrocytes. a, Peak-level summaries of diagnosis-associated differentially accessible peaks (DAPs) in PFC basal astrocytes, including a MA plot of ranked Log2Mean, a heatmap of DAPs across diagnosis, and genomic annotation summaries for all peaks, DS-enriched peaks, and DSAD-enriched peaks. b, Comparison of promoter accessibility changes and RNAexpression changes for DEGs in PFC basal astrocytes. Each point represents a gene and is colored according to the concordance between promoter accessibility and transcript-level changes, as indicated in the figure. c, rGREAT enrichment analyses for DS-enriched and DSAD-enriched peaks. Dot plots show selected Gene Ontology biological process terms, with dot size and color representing the enrichment statistics indicated in the figure.

**Extended Data Fig. 8.**
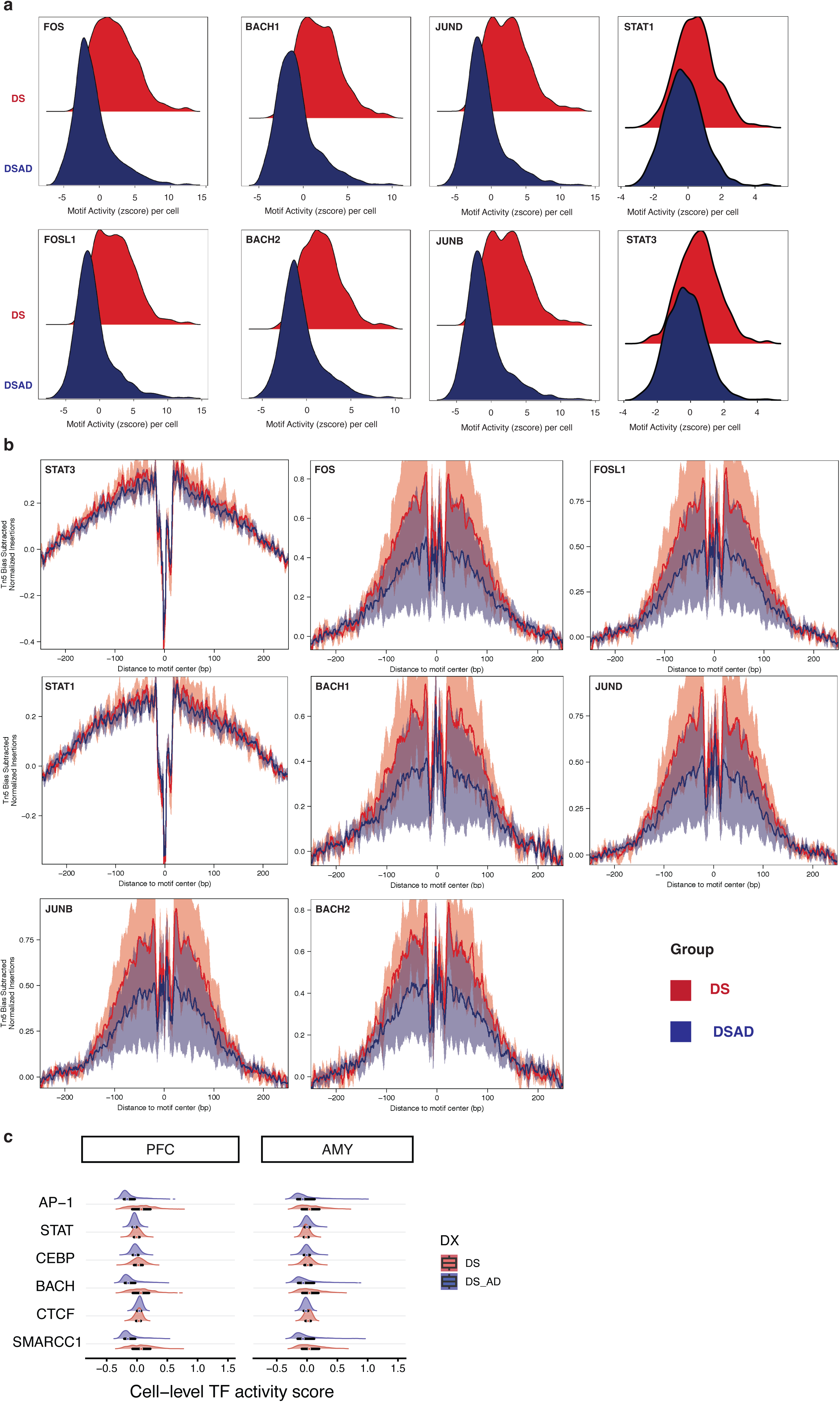
Additional chromVAR and footprinting analyses in PFC basal astrocytes. a, Density plots of chromVAR motif deviation scores for representative transcription factors in PFC basal astrocytes, comparing DS and DSAD including FOS, FOSL1, JUNB, JUND, BACH1, BACH2, STAT1 and STAT3. b, Motif-footprinting profiles for representative transcription factors, including STAT1, STAT3, FOS, FOSL1, BACH1, BACH2, JUND, and JUNB, comparing aggregated signal between DS and DSAD. Lines and shaded regions are displayed as indicated in the figure. c, Cell-level transcription factor activity score distributions for selected transcription factor families, shown separately for PFC and AMY with DS and DSAD displayed as labeled.

**Extended Data Fig. 9.**
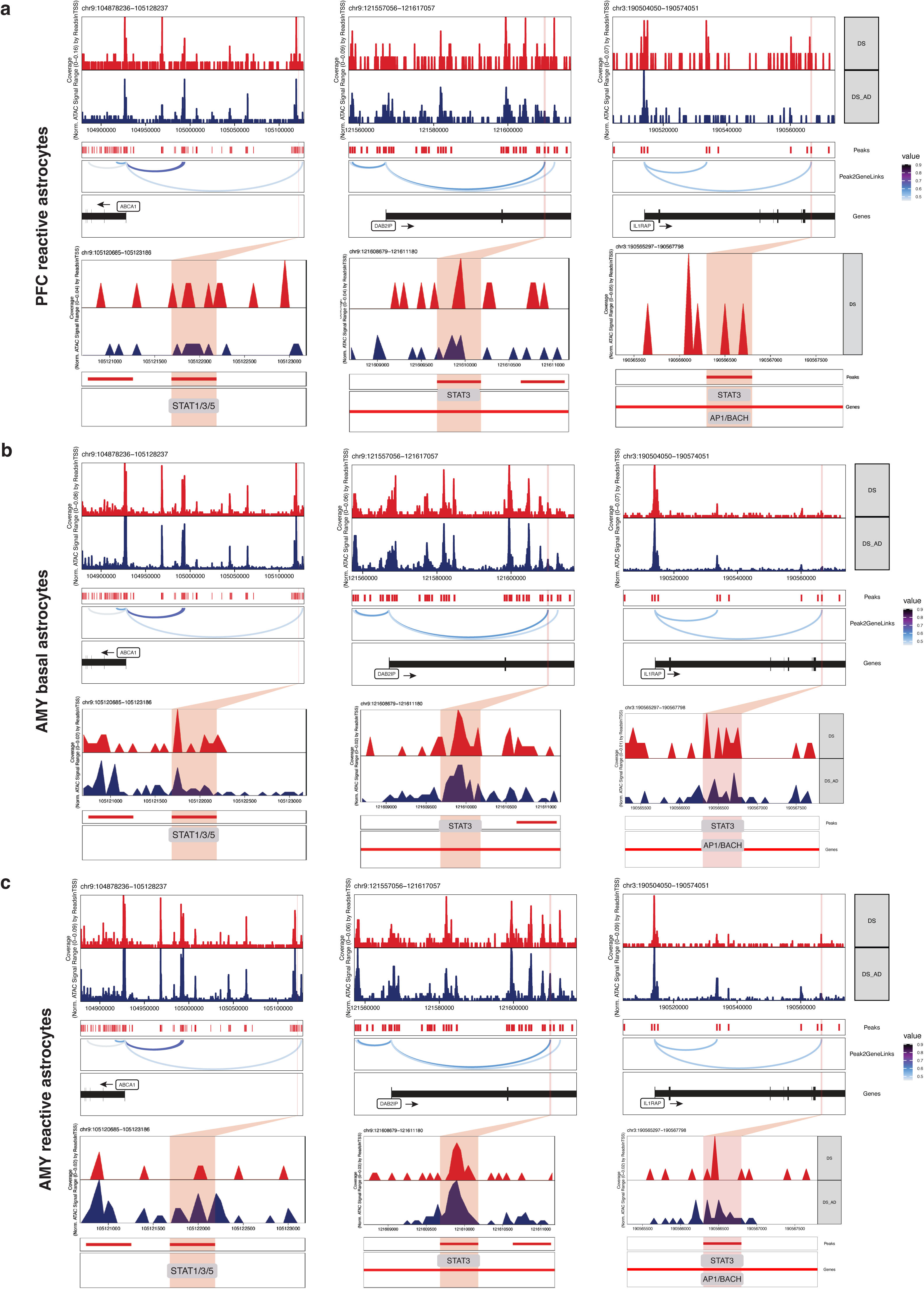
Comparison of candidate cis-regulatory loci across astrocyte states and brain regions. a-c, Genome-browser views of the ABCA1, DAB2IP, and IL1RAP loci in DS and DSAD astrocytes, shown for PFC reactive astrocytes (a), AMY basal astrocytes (b), and AMY reactive astrocytes (c). Tracks display aggregated accessibility signal together with peaks, peak-to-gene links where shown, gene annotations, and motif annotations for the highlighted candidate regulatory elements.

**Supplementary Fig. 1.**
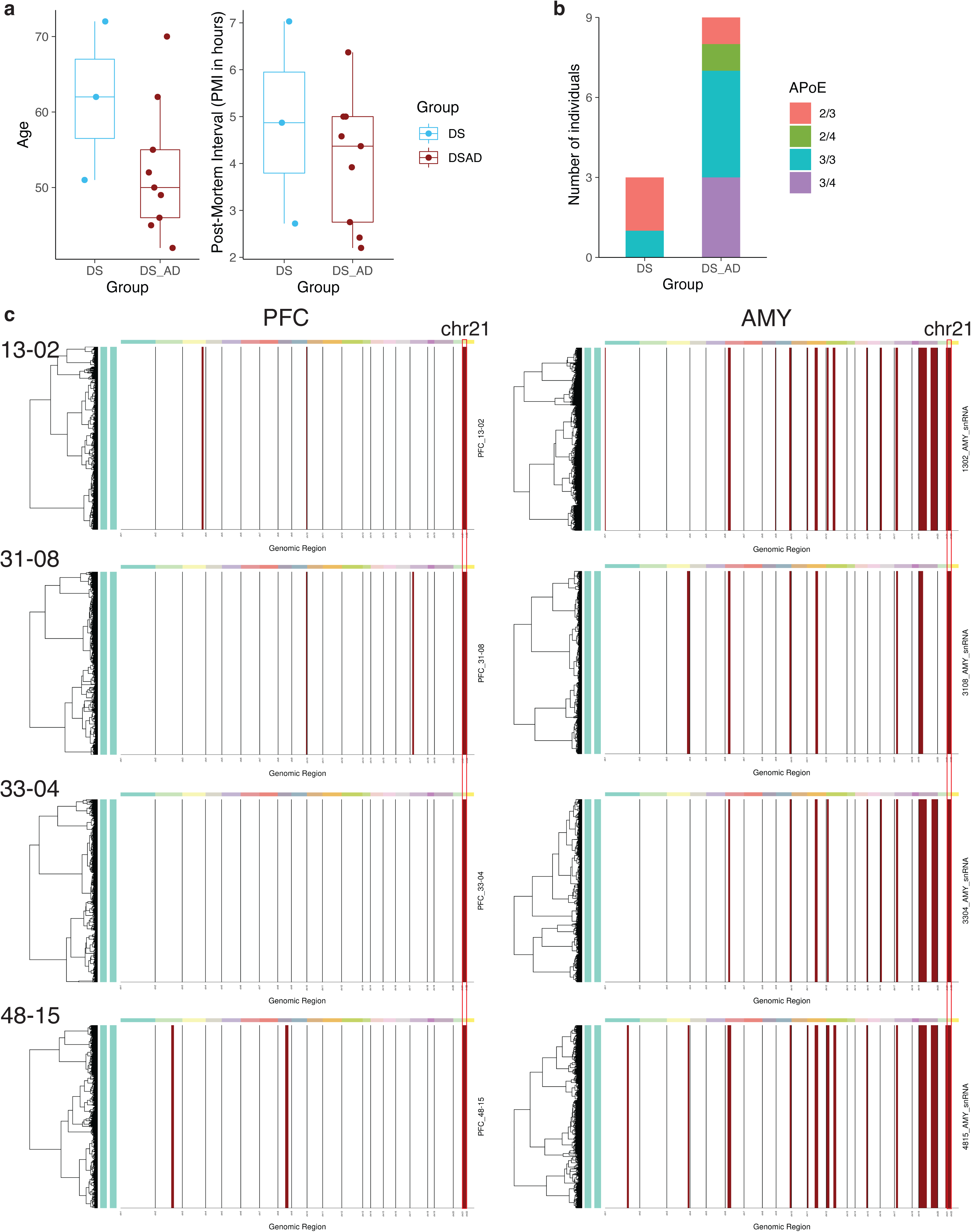
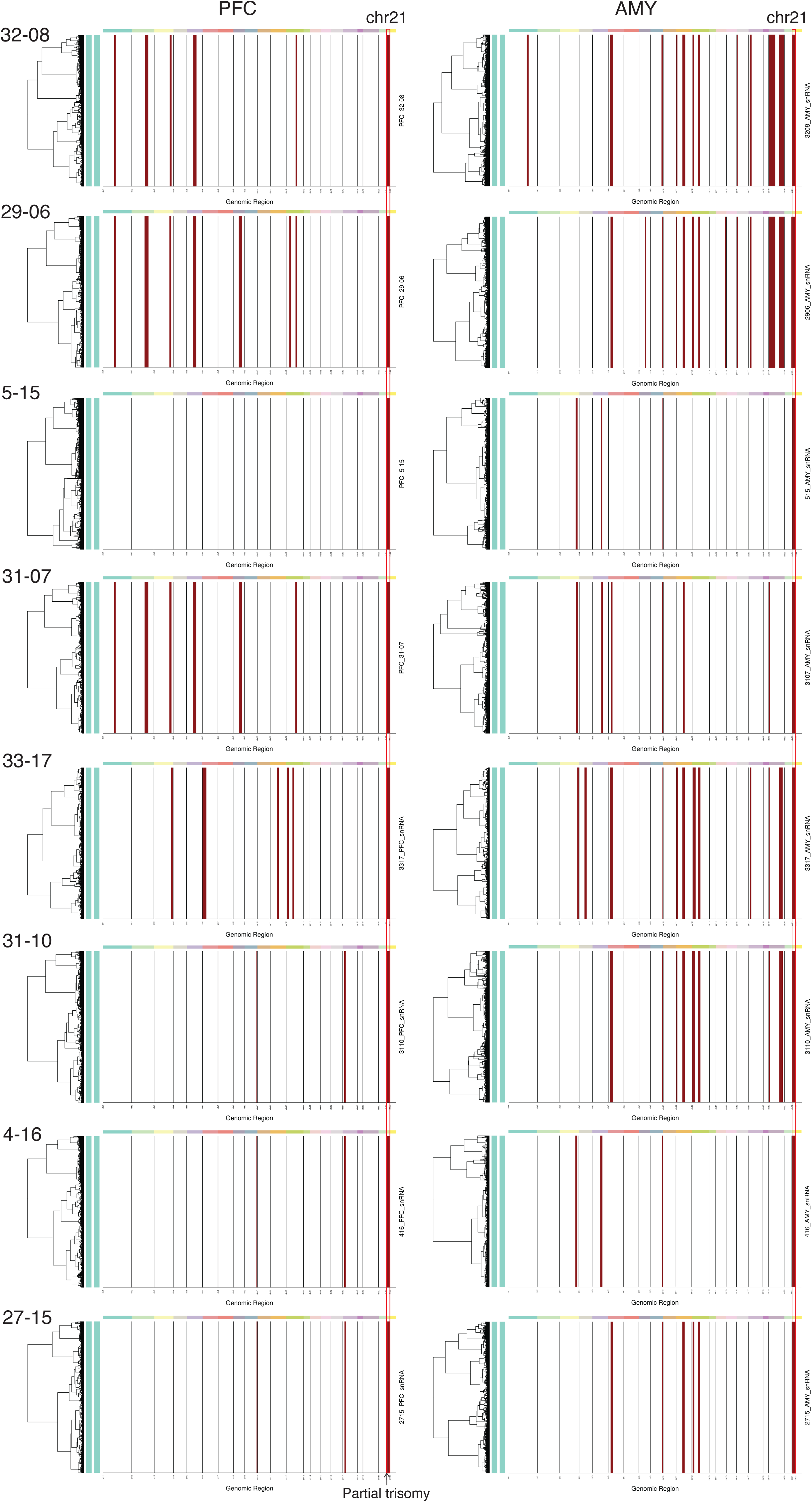
Cohort-level comparison of clinical variables and inferred chromosome 21 dosage across the DS and DSAD groups. a, Comparison of donor age and post-mortem interval (PMI) between the DS and DSAD groups. Boxplots summarize the group distributions and points indicate individual donors. b, APOE genotype distribution across donors in the DS and DSAD groups. Bar heights indicate the number of individuals. c, Inferred copy-number profiles from snRNA-seq data for each donor-region sample from the PFC and AMY, indicating chromosome 21 dosage.

